# Immune checkpoint expression on HIV-specific CD4+ T cells and response to their blockade are dependent on lineage and function

**DOI:** 10.1101/2021.09.23.461484

**Authors:** Elsa Brunet-Ratnasingham, Antigoni Morou, Mathieu Dubé, Julia Niessl, Amy E. Baxter, Olivier Tastet, Nathalie Brassard, Gloria Ortega-Delgado, Roxanne Charlebois, Gordon J. Freeman, Cécile Tremblay, Jean-Pierre Routy, Daniel E. Kaufmann

## Abstract

**Background:** Antigen-specific T cell impairment is observed in chronic infections. CD4+ T cells are diverse in phenotype and function; how their different lineages are impacted by inhibitory immune checkpoints (IC) is unknown.

**Methods:** We examined IC expression and function in HIV-specific CD4+ T cells of viremic individuals prior to ART initiation and persons with spontaneous or therapy-induced viral suppression. We investigated IC patterns associated with exhaustion-related transcription factors and chemokine receptors using cytokine-independent activation-induced marker assays. We determined effector functions representative of T_FH_, T_H_1 and T_H_17/T_H_22 using ultra-sensitive RNA flow cytometric fluorescence in situ hybridization (FISH), and their response to IC blockade.

**Findings:** The dysfunction-related transcription factor TOX was elevated in HIV-specific CD4+ T cells of viremic patients, and its expression was associated with lineage differentiation. We observed a hierarchy of PD-1, TIGIT and CD200 expression associated with both infection status and effector profile. *In vitro* responsiveness to PD-L1 blockade varied with defined CD4+ T cell functions rather than IC expression levels: frequencies of cells with T_H_1- and T_H_17/T_H_22-, but not T_FH_-related functions, increased. Response to PD-L1 blockade was strongest in viremic participants and reduced after ART initiation.

**Interpretation:** Our data highlight a polarization-specific regulation of IC expression and differing sensitivities of antigen-specific Thelper subsets to PD-1-mediated inhibition. This heterogeneity may direct ICB efficacy on CD4+ T cells in HIV infection.

**Funding:** This work was supported by the National Institutes of Health, the Canadian Institutes for Health Research, the Canada Foundation for Innovation and the Fonds de Recherche du Québec-Santé.

**Research in Context:** *Evidence before this study:* Combination antiretroviral therapy (ART) is highly effective in controlling HIV but requires life-long medication due to the latent viral reservoir, and does not restore suppressive immune responses. In particular, there is no generation of effective HIV-specific T cell responses, which are thought to play an important role in controlling HIV in the rare individuals who can spontaneously control the virus. Inhibitory immune checkpoints (IC) such as PD-1 contribute to T cell dysfunction and failure to control viral infections, including HIV, and IC blockade (ICB) represents a potential adjuvant to ART through restoration of T cell functions. While most studies have focused on CD8+ T cells, increasing evidence shows that the remarkable impact of ICB therapy in a subset of cancer patients is enhanced by functional CD4+ T cell help, which can be directly affected by ICB. While effective virus-specific CD4+ T cell responses are also thought to be important for immune control of HIV, these cells are highly heterogenous. How IC expression and function differs across CD4+ T cell lineages and the consequences of this diversity for IC blockade (ICB) strategies are still poorly understood.

*Added value of this study:* To compare various stages of immune dysfunction, we examined people living with HIV (PLWH) with different levels of viral control pre-ART (including elite controllers who spontaneously control virus) and followed a cohort longitudinally post-ART. We used a panel of assays to characterize HIV-specific CD4+ T cell subsets, including activation-induced marker (AIM) assays and flow cytometric detection of mRNAs coding for a wider variety of HIV-specific CD4+ T cell functions than what is detected by standard procedures. Our experiments indicate a hierarchy of IC (PD-1, TIGIT, CD200) expression on blood HIV-specific CD4+ T cells that depends not only on the person’s infection status but also on expression of lineage differentiation markers and effector functions representative of CD4+ T cell subsets critical for antiviral responses (T_FH_, T_H_1 and T_H_17/T_H_22 cells). This hierarchy was also present in the putatively functional cells of elite controllers. We characterized the expression of the dysfunction-related transcription factor TOX, and saw that its association with the key IC PD-1 in the setting of viremia varied across CD4+ T cell polarizations. Response to blockade of the PD-1 pathway resulted in increased antiviral and mucosal-protective functions, but did not affect T_FH_-related functions. Response to ICB was most prominent in viremic patients, and subdued but not fully abrogated in the setting of viral suppression.

*Implications of all available evidence:* These results highlight a previously unrecognized impact of ICB on mucosal immunity-related CD4+ functions, which are known to be depleted upon HIV infection and not restored by ART, and strong links between IC expression patterns and HIV-specific CD4+ T cell differentiation. The impact of ICB on CD4+ T cells in HIV infection has primarily been studied in the context of viral reservoir reactivation, which are preferentially harbored in IC+ cells. Our work emphasizes the importance of considering the differentiation profile of the virus-specific CD4+ T cells in studies of ICB blockade, as it may direct ICB efficacy in HIV infection. This data may also have implications for CD4+ T cell help in other infectious and non-infectious chronic human diseases.

## Introduction

CD4+ T helper (T_H_) cells orchestrate the immune responses against pathogens (1, 2) and defects in T helper responses contribute to lack of viral immune control in HIV infection. This diverse cell population polarizes towards lineages characterized by expression of chemokine receptors and transcription factors (TF), and produce distinct sets of cytokines (3). Beyond the prototypical antiviral T_H_1 subset, HIV-specific CD4+ T cells also include mucosal-related T_H_17/T_H_22 and B-cell helper T_FH_, the proportions of which are differentially related to spontaneous viral control (4). In chronic infections such as HIV, sustained antigenic exposure and inflammation alter both CD4+ and CD8+ T cell function. CD8+ T cell exhaustion follows a gradient enforced by epigenetic remodeling with limited reversibility (5, 6). TOX is a central transcription factor (TF) involved in the development and maintenance of exhausted CD8+ T cells in mice (7) and humans (8), although its role in human CD8+ T cells is not limited to exhaustion (9). Dysfunctional CD4+ T cells differ from exhausted CD8+ T cells in that they present prominent features of altered differentiation: loss of antiviral and mucosal-protective functions and a skewing towards a T follicular helper (T_FH_)-like profile (4). Little is known about TFs implicated in CD4+ T cells dysfunction, although some, increased in mice models, overlap with exhaustion-related TF (10). Another commonality between dysfunctional CD4+ and CD8+ T cells is the upregulation of inhibitory immune checkpoints (IC) (11), however with some notable differences in the IC hierarchy between the two subsets (10, 12-14). IC have dual roles as physiologic regulators of T cell activation and mediators of exhaustion (15). PD-1 is the best characterized IC contributing to both HIV-specific CD4+ and CD8+ T cell dysfunction (16), and correlates with disease progression (12, 14) and loss of antiviral function (16).

Immune checkpoint blockade (ICB) can partially rescue CD8+ T cell exhaustion, in particular blockade of the PD-1 signaling pathway. A population of mildly exhausted CD8+ T cells, called “progenitor exhausted”, with stem-like properties and intermediate levels of PD-1, respond to ICB(17-19), while terminally exhausted CD8+ T cells, with high PD-1 and Tim3+ expression, have poor response to ICB (19). Responsiveness of CD4+ T cells to ICB is understudied due to their heterogeneity and the paucity of tools to identify them in an antigen-specific manner. Although PD-1’s effect on CD4+ T cell function *in vivo* was classically described as IL-2 inhibition (20, 21), studies in animal and human chronic infections support broader ramifications. PD-1 blockade enhanced IFNγ+ T_H_1-responses specific to *Mycobacterium tuberculosis* in murine models (22) and patients undergoing ICB for cancer therapy (23), and moderately increased IFNγ secretion by SIV-specific CD4+ T in non-human primates (24). These primates had replenished T_H_17 in the gut and improved gut integrity, which may explain their improved survival through reduced immune hyperactivation (25, 26). *In vitro*, PD-1 blockade enhanced HIV-specific CD4+ T cell proliferation as well as IFNγ, IL-2, IL-13 and IL-21 production (14). However, the links between IC expression among the heterogeneous polarizations of CD4+ T cells and the impact of ICB on various effector functions of these cells are still lacking.

Here, we define the relationships between dysfunction-related characteristics and the lineages of HIV-specific CD4+ T cells across disease and treatment status. We pinpoint a previously underappreciated heterogeneity across types of CD4+ T cells in both their IC and exhaustion-related TF expression patterns, as well as their capacity to respond to PD-1 blockade. Greater understanding of ICB’s impact on CD4+ T cells can foster new therapeutic uses for immunotherapeutic interventions.

## Materials and Methods

### Study Design

Leukaphereses were obtained from study participants at the Montreal General Hospital, Montreal, Canada and at the Centre Hospitalier de l’Université de Montréal (CHUM) in Montreal, Canada. The study was approved by the respective IRBs (IRB CHUM: 17.335) and participants gave written informed consent prior to enrollment. Samples were collected between 2013 and 2019 as part of a multicentric study (MP-37-2018-4029). Subject characteristics are summarized in Supplementary Table 1. Chronic Progressors (CP) had plasma viral loads of at least 5000 copies/ml and were infected and infected/off treatment for at least 3 months at the time of collection of the “Pre-ART” sample. Longitudinal “Post-ART” samples were collected in these same subjects, after at least 3 months on ART and undetectable viral loads. Elite controllers (EC) had spontaneously controlled viremia (< 40 viral copies/ml) in the absence of ART. PBMCs were isolated by the Ficoll density gradient method and stored in gas phase of a liquid nitrogen tank in 90%FBS with 10% DMSO.

### Antibodies

All antibodies are listed in Supplementary Tables 2-5. Antibodies are monoclonal and raised in mice. All antibodies were validated by manufacturer and titrated with biological and/or isotype controls.

### Activation-induced marker (AIM) assay

As previously described (4), cryopreserved peripheral blood mononuclear cells (PBMCs) were thawed and rested in cell culture media (RPMI supplemented with 10% Human AB serum and PenStrep – 50 U/ml of penicillin and 50 µg/ml of streptomycin) at 37°C for 3 hours at a density of 10M/ml in 24-well plates. 15 minutes prior to stimulation, CD40 blocking antibody (clone HB14, Miltenyi, cat #: 130-094-133) was added to each well at 0.5 µg/ml, as well as antibodies staining CXCR5, CXCR3 and CCR6. Cells were either left unstimulated or stimulated with overlapping peptide pools of HIV Gag (JPT, PM-HIV-Gag ULTRA), at a final concentration of 0.5 µg/ml/peptide. Alternatively, 1µg/ml of Staphylococcal Enterotoxin B (SEB, Toxin Technology) was used to stimulated the cells as a positive control. Cells were stimulated for 9 hours, collected, washed and stained with LIVE/DEAD™ Fixable Aqua Dead Cell Stain Kit (20 mins, 4°C; Thermofisher, #L34965). After washing, cells were incubated with FcR block (10mins, 4°C; Miltenyi) then stained with a cocktail of surface markers (30 mins, 4°C; See panel in Supplementary Table 2). Washed cells were then fixed with 2% paraformaldehyde (PFA) for 20 mins at RT, then washed and resuspended in PBS-2% FBS for flow acquisition on a 5-laser LSRII (BD BioSciences). For experiments with intranuclear transcription factor staining, fixation and permeabilisation were done using eBioscience™ Foxp3 / Transcription Factor Staining Buffer Set (cat#: 00-5523-00) following kit instructions: surface-stained cells were fixed with 1x Fixation/Permeabilisation for 30 min at RT in the dark, then washed and resuspended in 1X Permeabilization buffer with intranuclear antibody cocktail for 1h at RT in the dark. Analysis was performed using FlowJo (Treestar, V10). Gates were set on the unstimulated controls.

### Combined cytokine/chemokine mRNA-Flow-FISH and protein staining assays

As previously described(4), PBMCs were thawed and rested for 2-3 hours in 48-well plates at 5M in 0.5ml in cell culture medium. 15 minutes prior to stimulation, a PD-L1 blocking antibody (29E.2A3(27)) or an isotypic control (IgG2b, clone MPC-11, BioXcell, # BE0086) at a concentration of 10 µg/ml were added into culture, along with antibodies staining CXCR5, CXCR3 and CCR6. PBMCs were then either left unstimulated or were stimulated with an HIV Gag peptide pool (JPT) or SEB for 12 hours. After incubation, cells were stained with Fixable Viability Dye eFluor™ 506 (20 min, 4°C; eBioscience, # 65-0866-14) before labeling of surface markers with surface antibodies (30 min, 4°C; See panel in Supplementary Table 3). Samples were next subjected to the PrimeFlow RNA® assay (ThermoFisher) for specific mRNA detection in a 96-well plate as per manufacturer’s instructions. All buffers and fixation reagents were provided with the kit, with the exception of flow cytometry staining buffer (PBS - 2% FBS). Briefly, after fixation and permeabilization, cytokine/chemokine mRNAs were labelled with one of five combinations of probes as listed in Supplementary Table 4. The probes were each diluted 1:20 in probe diluent and hybridized to the target mRNA for 2 hr at 40°C. Samples were washed to remove excess probes and stored overnight in the presence of RNAse inhibitor 1X (RNAsin). Signal amplification was achieved by sequential 1.5 hr incubations at 40°C with the pre-amplification and amplification mixes. Amplified mRNA was labelled with fluorescently-tagged probes for 1 hr at 40°C. Samples were acquired on a BD LSRFortessa™. Analysis was performed using FlowJo (Treestar, V10). Gates were set on unstimulated controls (see Fig S3b). Net frequencies of HIV-specific responses were calculated by subtracting the background expression in the absence of exogenous stimulation from the value measured after Gag antigen stimulation. HIV-specific responses were considered positive when the frequency obtained with Gag stimulation was at least twice that obtained in the absence of exogenous stimulation. Responses not meeting this criterion were characterized as negative.

### Delayed Intracellular cytokine staining

As previously described(28), thawed, rested PBMCs were either left unstimulated or were stimulated with an HIV Gag peptide pool (JPT) or SEB. After a 9h stimulation, 1.25 µg/ml of brefeldin A (BD GolgiPlug) was added to culture and cells were further incubated for 12 hours. Cells were collected, washed and stained with AquaVivid Viability dye (20 mins, 4°C). After washing, cells were incubated with FcR block (10mins, 4°C) then stained for cocktail of surface markers (30 mins, 4°C; see supplementary table 5 for panel). Cells were washed and fixed with Fixation Solution (eBioscience, #88-8824-00) for 15 mins at RT, following which they were washed and stained for intracellular proteins with 1X Permeabilization Buffer (eBioscience, #00-8333-56) (30 mins, 4°C). Cells were washed once more with 1X Permeabilization buffer, then with PBS – 2%FBS and acquired on the BD LSRFortessa. Analysis was performed using FlowJo (Treestar, V10). Gates were set on unstimulated controls.

### qRT PCR analysis of HIV-specific CD4+ T cells

These data were collected in a previously published study (4). Briefly, AIM assay was conducted as previously explained and CD69+CD40L+CD4+ T cells were live-sorted on a FACS Aria cell sorter (BD BioSciences) equipped for handling of biohazardous material, operated at 70 pounds per square inch with a 70-um nozzle (for gating strategy, please refer to (4). 5000 cells were collected directly into RLT lysis buffer (Qiagen) and vigorously vortexed before flash-freezing. Total RNA was purified using the RNeasy Plus Micro Kit (Qiagen). cDNA was synthesized using all RNA available (or 1-5 ng) with the High-Capacity Reverse Transcription Kit with RNase Inhibitor (Life Technologies) (250 C for 10 min, 370 C for 120 min, 850 C for 5 min). cDNA equivalent to 1000 sorted cells was subjected to gene-specific preamplification using Taqman Preamp MasterMix (Applied Biosystems) and 96 pooled TaqMan Assays (Applied Biosystems – for full panels, please refer to (4) at final concentration 0.2X (95°C for 10 min, followed by 16 cycles of 95°C for 15 s and 60°C for 4 min). The preamplified cDNA was diluted 5-fold in DNA suspension buffer (Teknova) and was mixed with TaqMan Universal PCR Master mix (Life Technologies) and 20X GE sample loading reagent (Fluidigm). 20X Taqman assays were diluted 1:1 with 2X assay loading buffer (Fluidigm). Taqman assays mixtures were loaded onto a primed 96.96 Dynamic Array chip (Fluidigm). The chip was loaded into the IFC Controller, where each sample was mixed with each assay in every possible combination. The chip was transferred in a Biomark (Fluidigm) for real-time PCR amplification and fluorescence acquisition using single probe (FAM-MGB, reference: ROX) settings and the default hot-start protocol with 40 cycles. Cycle thresholds (Ct) were calculated using the Fluidigm BioMark software.

Analysis of the qRT-PCR data obtained on the microfluidic platform was carried out using GenEx software (MultiD Analyses, URL: http://www.multid.se). Five endogenous control genes were included in the Fluidigm run and the stability of endogenous control genes across all experimental samples was evaluated applying the NormFinder algorithm50 in GenEx. The mean expression of the most stable endogenous control genes was used for normalization and calculation of -ΔCt values. Principal component analysis and biplots were created using the prcomp and fviz_pca_biplot functions in R programming language.

### Statistical analyses

Statistical analyses were performed with Prism v6.0 (GraphPad) using non-parametric tests. The type of statistical test is specified in the figure legends. Permutation test (10 000 permutations) was calculated using the SPICE software (https://niaid.github.io/spice/). Statistical tests were considered two-sided and p<0.05 was considered significant. The heatmap, dendrogram and PCA were generated using the fold change between the net value of the frequency of a cytokine mRNA detected with PD-L1 blockade over that seen for the same cytokine with the isotypic control. The prcomp function was used for the PCA, and the ggfortify and pheatmap packages were used for the dendrogram and heatmap, respectively.

## Results

### Exhaustion-related transcription factor TOX correlates with PD-1 expression

To explore dysfunction among the heterogeneous T helper (T_H)_ populations, we compared dysfunctional HIV-specific CD4+ T cells from viremic chronic progressors with high viral burden prior to ART (CP; VL > 5000 viral RNA copies/ml) to the relatively functional HIV-specific CD4+ T cells from elite controllers who spontaneously suppress virus (EC; VL < 40 copies/ml) (patient characteristics in Supplemental Table 1)(4). Upregulation of activation-induced markers (AIM) following peptide stimulation allows the capture of a broader antigen-specific CD4+ T cell population than cytokine-based techniques (29). We stained for co-expression of CD69 and CD40L, an activation-induced co-signaling molecule expressed on multiple polarizations but low on bystander activated cells (4, 29), after a 9-hour *ex vivo* stimulation with a peptide pool of HIV’s immunodominant antigen, Gag (Fig 1a and Fig S1a). Both cohorts had similar frequencies of AIM+ HIV-specific CD4+ T cells (Fig 1b), and PD-1 expression was higher on HIV-specific CD4+ T cells from CP than EC (Fig 1c, and Fig S1b), as previously reported (12, 14).

**Figure 1.**
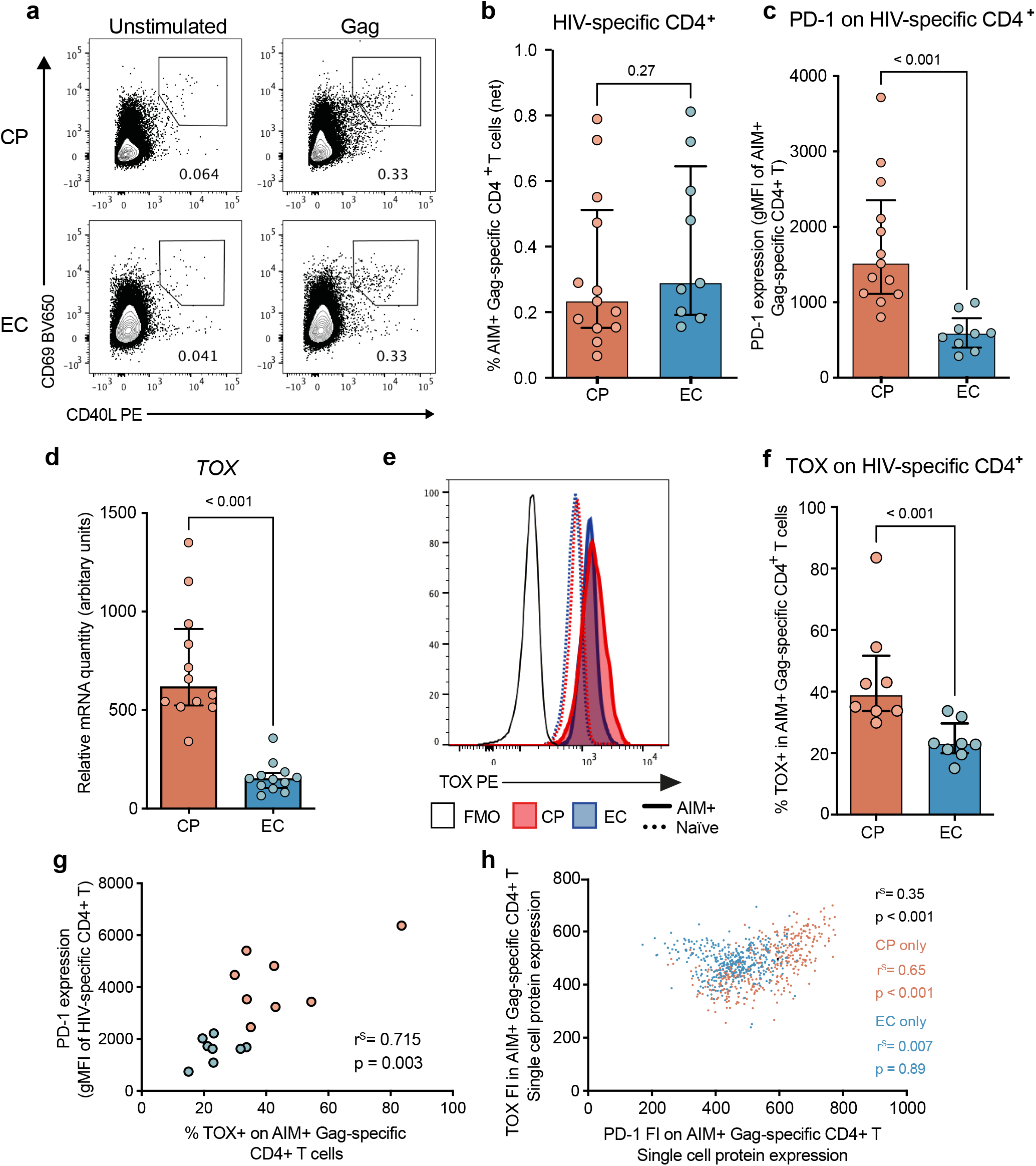
Increased expression of exhaustion-related transcription factors in HIV-specific CD4+ T cells of CP compared to EC. **a**) Representative flow cytometry plots and **b**) cumulative data of AIM+ Gag-specific CD4+ T cells detection via upregulation of the activation-induced markers (AIM) CD69 and CD40L in a CP (top) and an EC (bottom) 9 hours after stimulation with a HIV Gag peptide pool. **c**) Comparison of PD-1 expression on AIM+ HIV-specific CD4+ T cells in the CP (orange) and EC (blue) cohorts. Representative flow cytometry plot of TCF-1 and TOX staining on total CD4+ T cell (top) or Gag-specific AIM+ CD4+ T cells (bottom). **d**) Relative *Tox* mRNA expression among sorted Gag-specific CD4+ T cells of CP (red) or EC (blue), as captured by high-throughput RT-PCR (Fluidigm® - for details, please see (4). **e**) Representative example of TOX expression in AIM+ HIV-specific CD4+ T cells (shaded) or unstimulated naïve CD4+T cells (dotted line) of both cohorts. Black line is FMO control. **f**) Cumulative data of the frequency of TOX+ cells among AIM+ Gag-specific CD4+ T cells (right) of CP (blue) or EC (red), using gating strategy depicted in A. **g**) Correlation between the frequency of TOX and PD-1 expression level among AIM+ Gag-specific CD4+ T cells. **h**) Correlation between the single cell expression (as captured by flow cytometry – FI = fluorescence intensity) of TOX and PD-1. Correlation done using 100 cells per patient for 4 CP and 4 EC. bc) n = 13 CP and 9 EC. d) n = 9 CP and 9 EC. fg) n = 8 CP & 8 EC. bcdf) Stats = Mann-Whitney test. GH) Spearman correlation.

We considered the exhaustion-related TF TOX, given its association with T cell exhaustion(7), and TFs reported in mouse dysfunctional CD4+ T cells (10): Blimp-1 (Prdm1), Helios, Nfatc1, Batf, Eomes and Tbet. Our previously published high-throughput qRT-PCR assay data on sorted HIV-specific CD4+ T cells from CP or EC (4) showed increased *TOX* (Fig 1d*), IKZF2* (HELIOS), *NFATC1*, and *PRDM1* (BLIMP-1) (Fig S1c) in HIV-specific CD4+ T cells of CP compared to EC, while *BATF* and *TBX21* were higher in EC. *EOMES* was similar in both cohorts.

We next performed intra-nuclear protein staining for the TF (which were largely increased in CP compared to EC) in HIV-specific CD4+ T cells using the 9h AIM assay (Fig S1d). HELIOS was undetectable in the AIM+ population, whereas both NFATc1 and TOX were increased in the CP compared to the EC. TOX was most differential based on varying viral loads (Fig S1d). As CD4+ T cell dysfunction and viral load are strongly associated in HIV infection (4), TOX was best candidate for assessment of dysfunction by flow cytometry.

We set the TOX+ gate on naïve CD4+ T cells (Fig 1e), and confirmed a greater frequency of TOX+ cells in AIM+ HIV-specific CD4+ T cells of the CP cohort compared to EC (Fig 1f). TOX correlated significantly with PD-1 expression at the patient level (Fig 1g) and at the single-cell level (Fig 1h). Of note, EC had a population of TOX+PD-1low cells not observed in CP (Fig 1g), losing the correlation between PD-1 and TOX single cell expression. These observations suggest TOX and PD-1 are increased jointly in the setting of dysfunction, perhaps from common upregulating signals.

### Differential PD-1 expression in polarized HIV-specific CD4+ T cells

Among AIM+ HIV-specific CD4+ T cells, we characterized three polarizations based on chemokine-receptors expression: CXCR3, CCR6 and CXCR5, enriched on antiviral T_H_1, mucosal-related T_H_17/T_H_22 and B-cell helper T_FH_, respectively (Fig 2a). Proportions were comparable between CP and EC, with the exception of a decreased CCR6+ fraction in CP (Fig S2a), as previously reported(4). TOX expression varied among polarizations and the hierarchy was not maintained between both cohorts: while in CP, the CXCR3+ polarization had significantly greater TOX levels than CCR6+ and CXCR5+, in EC TOX levels were greater in CCR6+ than CXCR5+, and intermediate in CXCR3+ (Fig 2b).

**Figure 2.**
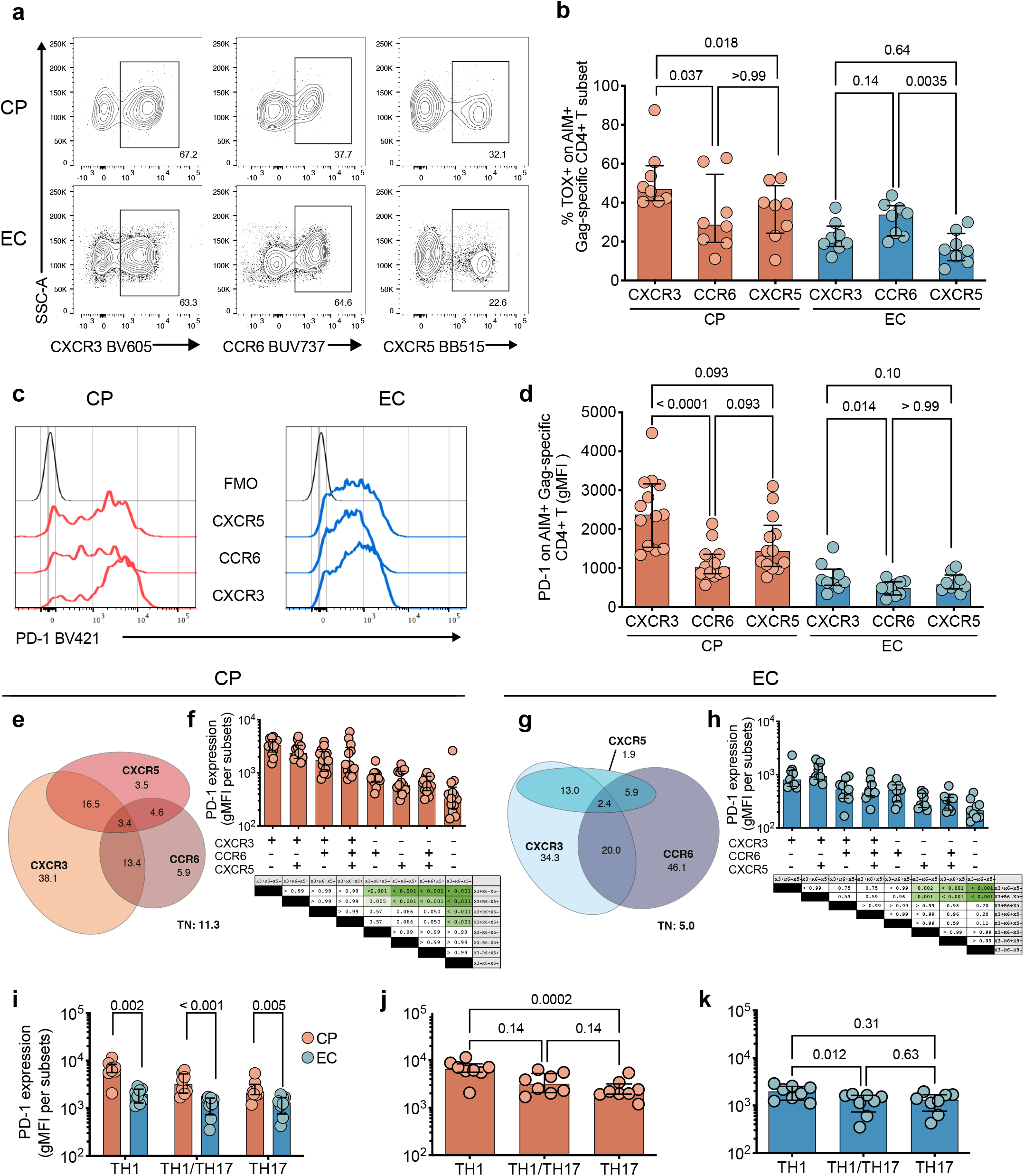
PD-1 expression on HIV-specific CD4+ T cells depends on their polarization. **a**) Representative flow cytometry plots of expression of the chemokine receptors CXCR3, CCR6 and CXCR5 on AIM+ Gag-specific CD4+ T cells of a CP (top) and an EC (bottom). **b**) Expression of TOX among chemokine-receptor-identified polarizations of Gag-specific CD4+ T cells among CP (red) or EC (blue). **c**) Representative example and **d**) cumulative data of PD-1 expression among chemokine-receptor-identified polarizations of Gag-specific CD4+ T cells. Euler graphs of co-expression for CXCR3, CCR6 and CXCR5 on AIM+ Gag-specific CD4+ T cells in **e**) CP or **g**) EC. Values represent median frequencies of subsets within total AIM+ Gag-specific CD4+ T cells. TN: triple negative for all three chemokine receptors. PD-1 expression in AIM+ HIV-specific CD4+ T cells subsets, as identified by chemokine co-expression patterns in **f**) CP or **h**) EC. Statistics appear in tables below, with p values < 0.05 highlighted in green. PD-1 expression on CD4+ T cell subsets identified by chemokine receptor and master transcription factors, **i**) in CP vs EC; between polarizations of **j**) CP or **k**) EC. In D-H n= 13 CP and 9 EC. I-J n = 8 CP and 8 EC. Columns correspond to median values with interquartile range. dfhjk) Friedman test with Dunn’s post-test. i) Mann-Whitney test. Geo MFI: Geometric Mean of Fluorescence Intensity.

PD-1 expression also varied among polarizations, mimicking TOX’s pattern in CP: highest PD-1 was again observed on CXCR3+, while CXCR5+ and CCR6+ had comparable levels (Fig 2cd). In EC, the hierarchy of PD-1 was similar to that of CP, but contrasted with the hierarchy of TOX in EC: CXCR3+ cells had the greatest PD-1 levels, although only significantly greater when compared to CCR6+ cells. This is in line with the absence of correlation for EC between single-cell expression PD-1 and TOX, and further emphasizes that PD-1 and TOX expression are specifically linked in the context of dysfunction.

CXCR3, CCR6 and CXCR5 can be co-expressed in various patterns, in line with the plastic nature of T_H_ (Fig 2eg). PD-1 expression was highest on the CXCR3+ CCR6-subsets (Fig 2fh). We further examined co-expression of classical “master” TFs with chemokine receptors, identifying T_H_1 as CXCR3+ T-BET+EOMES+, T_H_17 as CCR6+ROR-γt+CXCR3- and T_H_1/T_H_17 as CCR6+ROR-γt+CXCR3+(Fig S2e) (30). T_FH_’s master regulator BCL-6 was largely undetectable in peripheral CD4+ T cells (Fig S2e), as previously reported (31). Among the AIM+ HIV-specific CD4+ T cells, the proportions of T_H_17 and T_H_1/T_H_17 were significantly higher in EC, but similar for T_H_1 (Fig S2f). PD-1 expression in CP always exceeded that in EC (Fig 2i) and, in both cohorts, TH1 cells had greater PD-1 expression than a CCR6+ polarization (Fig 2jk).

Thus, TOX and PD-1 expression follow similar patterns in the setting of dysfunction only ; however, the differing hierarchy of PD-1 expression among subsets of HIV-specific CD4+ T cells is observed both in CP and in EC.

### PD-1 levels differ according to HIV-specific CD4+ T cell functions

We further identified HIV-specific CD4+ T cell subsets by cytokine expression and cytotoxic functions. Flow cytometric RNA fluorescent *in situ* hybridization (RNA-Flow-FISH) assay can capture hard-to-detect cytokines transcribed by HIV-specific CD4+ T upon cognate antigen stimulation, with fluorescence intensity giving a semi-quantitative measurement of the number of RNA copies per cell (32). We examined eight cytokines plus granzyme B (GZMB) that spanned five functional categories: IFNγ and IL-2 for T_H_1-associated functions; GZMB for cytotoxic activity; IL-22 and IL-17F for mucosal-related T_H_17/T_H_22 functions; IL-21, CXCL13 and IL-4 for T_FH_-associated functions; and IL-10, a pleiotropic molecule with mostly inhibitory functions (Fig 3a, Fig S3a). CD69 served as a surrogate for recent activation to increase specificity for HIV antigen-induced cytokine mRNA (Fig S3b). HIV-specific CD4+ T cells producing *IL-4* and *IL-10* mRNA had low or undetectable frequencies in most participants and were not pursued (Fig S3bc). CPs had lower frequencies of *IL22* mRNA+ cells, with a trend for lower *IL17F* and *GZMB* mRNA+ cells (Fig 3b), and fewer mucosal-related cytokine transcripts per cell (Fig 3C). Conversely, CP exhibited increased frequencies of T_FH_-related cytokines *IL21* and *CXCL13* mRNA+ cells (Fig 3b). Polyfunctional cells were observed in both cohorts, with only the *GZMB+IL2+* populations rarely detected (Fig S3d). The other combinations followed the expected trends (Fig 3b): *GZMB* single-positive cells and all combinations of mucosal cytokines were greater in EC, whereas the CP had higher frequencies of T_FH_-related cytokines combinations. *IFNG* single-positive cells were increased in the response of CP, consistent with the reported loss of polyfunctionality in HIV-specific T_H_1 (33).

**Figure 3.**
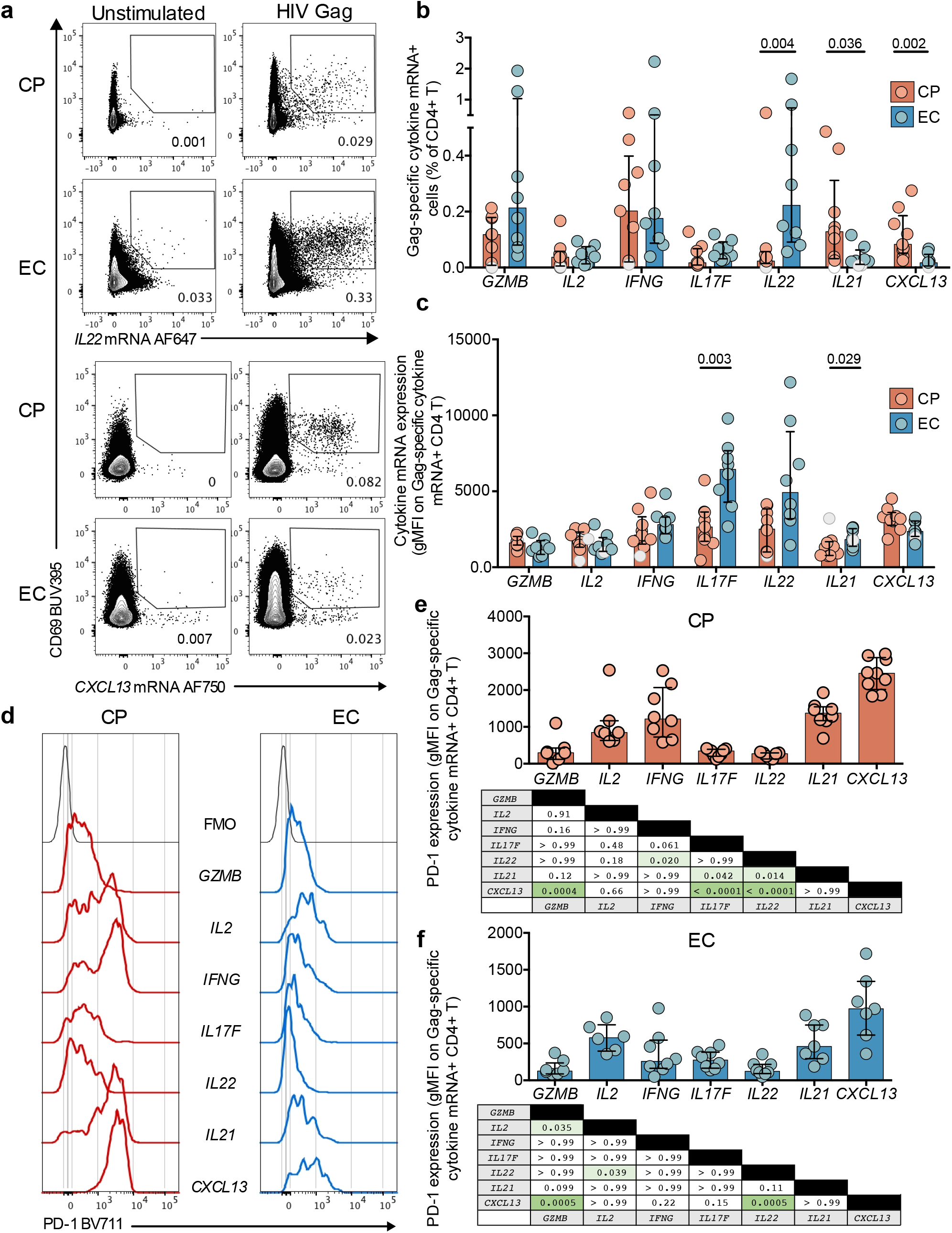
Heterogeneous PD-1 expression among cytokine-producing HIV-specific CD4+ T cells. **a**) Representative flow cytometry plots of IL22 mRNA and CXCL13 mRNA detection in a CP and an EC. Cumulative data of the **b**) net frequencies or **c**) geometric mean fluorescence intensity of Gag-specific cytokine mRNA+ CD4+ T cells in both cohorts. **d**) Representative examples and cumulative data of PD-1 expression on Gag-specific cytokine mRNA+ CD4+ T cells in **e**) CP or **f**) EC. Statistics appear in tables below, with p values < 0.05 highlighted in green. All mRNA data were acquired following a 12-hour stimulation with HIV Gag peptide pool. In bcef, n = 9 CP and 8 EC. In ef) only positive responses (at least 2 fold greater than unstimulated) are considered. Negative responses identified by grey shapes in bc. Bars represent medians with interquartile range. bc) Mann Whitney test. ef) Kruskall Wallis, with Dunn’s multiple comparisons tests.

Chemokine receptors expression among cytokine mRNA+ HIV-specific CD4+ T cells revealed complex associations between phenotype and function (Fig S3e). While CXCR3 was expressed on a large majority of *GZMB, IL2* and *IFNG* mRNA+ cells (Fig S4a-c), it was also present on most *IL21* and *CXCL13*+ mRNA cells. A minority of T_FH_-associated cytokine+ cells expressed CXCR5, with this proportion being smaller in CP. In contrast, almost all *IL22* or *IL17F* mRNA+ cells expressed CCR6.

Among defined T helper functions, PD-1’s hierarchy was similar to that observed on chemokine-receptor-identified polarizations: low on cells producing *GZMB* and mucosal-associated cytokines *IL22* and *IL17F*, and high on T_H_1 (*IFNG, IL2*) and T_FH_ (*IL21, CXCL13*) cytokine mRNA+ cells (Fig 3d-f). The hierarchy was similar in both cohorts, with the exception of particularly low PD-1 on *IFNG* mRNA+ cells in EC (Fig 3f).

These results demonstrate that HIV-specific CD4+ T cells can retain at least part of their functionality despite high PD-1 expression. Viremia leads to upregulation of this IC on functional cells, although the extent of its increase varies among T helper functions.

### Differential responsiveness of individual cytokines to PD-1 blockade

Given the hierarchical expression of PD-1 among CD4+ T cells of different functions, we speculated that responsiveness of these cells to blockade of PD-L1, the major ligand for PD-1 in PBMCs, would be heterogeneous as well. On cells from CP, PD-L1 blockade increased frequencies of HIV-specific cytokine mRNA+ CD4+ T cells for mucosal and antiviral functions (Fig 4ab), only slightly impacted *IL21*, and no effect on *CXCL13* mRNA+ cells. Blockade in EC had a globally smaller impact on HIV-specific CD4+ T cells (Fig 4cd), with increased frequencies only detected for *IL2, IFNG*, and *IL22*. Fold increases upon blockade of the 7 functions classified the participants relative to their cohort by unsupervised hierarchical clustering (Fig 4e), and in principal component analysis (PCA), clearly depicts disease status as the main source of variation (Fig 4f). CP were heterogeneous in which type of cytokine-producing cells were increased upon blockade, suggesting PD-L1 blockade does not result in a consistent profile of response even for one same antigen specificity. Many combinatory subpopulations increased in frequencies upon PD-L1 blockade in CP, with the notable exception of *CXCL13* mRNA+ cells (Fig 4ghi). EC had overall lower responses (Fig S4efg). For most constellations, there was at least a trend for greater response in CP than in EC in terms of fold change, although the spread in our relatively small cohorts did not allow to rank responsiveness to PD-L1 blockade among subsets.

**Figure 4.**
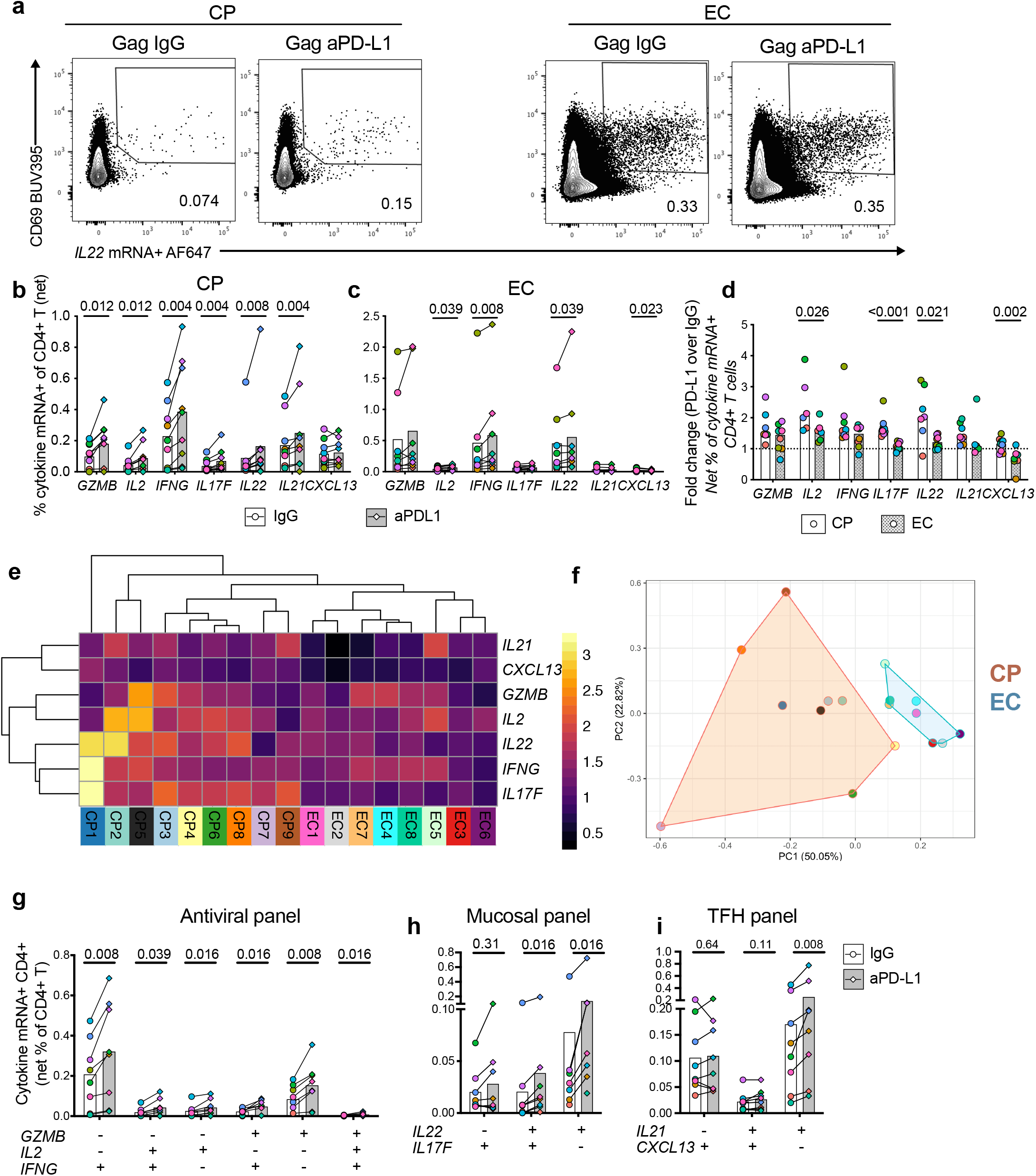
Differential responsiveness of individual HIV-specific CD4+ T cell cytokines to PD-1 blockade. **a**) Representative flow cytometry plots of *IL22* mRNA following Gag stimulation with PD-L1 blocking antibody (aPD-L1) or isotypic control (IgG) in a CP and an EC. Cumulative net frequency for all cytokine mRNA+ CD4+ T cells in **b**) CP and **c**) EC. **d**) Fold change in the net frequencies of cytokine mRNA+ HIV-specific CD4+ T cells upon PD-L1 blockade compared to isotypic control for both cohorts. CP in orange and EC in blue. **e**) Unsupervised hierarchical clustering analysis and heatmap of fold changes per cytokine across subjects, with warmer colors representing stronger fold changes. Bottom row corresponds to individual subject IDs. **f**) Principal component analysis (PCA) representation of CP (orange) and EC (blue) responses based on cytokine mRNA fold changes upon PD-L1 blockade. Length of lines to cytokines represent contribution to variance of each cytokine; angle of line represents their contribution to either PC1 and PC2. Red or blue shading regroups CP or EC, respectively. The numbers in parentheses are the percentage of variance explained by each principal component. Response of all cytokine mRNA combinations to PD-L1 blockade, upon Gag-stimulation among CP, for **g**) antiviral panel; **h**) mucosal panel or **i**) TFH panel. n = 9 CP and 8 EC. Columns correspond to median values with interquartile range. bcghi) Wilcoxon test. d) Mann-Whitney test. Each donor within a cohort has been separately color coded.

To study whether the effects observed on mRNA translated to protein, we performed delayed intracellular cytokine staining (d-ICS) following Gag stimulation. Extended stimulation before the addition of brefeldin A allows to capture the expression of both cytokines produced early, like IL-2 and IFNγ, as well as molecules induced later, namely CXCL13 and IL-21 (28). IL-17F and IL-22 were not detectable (Fig S4ab). Cytokine protein and cytokine mRNA expression correlated significantly for the IFNγ, IL-2 and CXCL13, but not for IL-21 (Fig S4c). Frequencies of IFNγ+ and IL2+ HIV-specific CD4+ T cells were increased upon blockade, but not CXCL13+ cells (Fig S4d), reflecting the lack of response to anti-PD-L1 seen at the mRNA level.

These data demonstrate a heterogeneous capacity of functionally-distinct HIV-specific CD4+ T cells to respond to PD-L1 blockade at the transcriptional and translational level, including for polyfunctional cells.

### IC co-expression on HIV-specific CD4+ T cells is lineage- and function-specific

To understand the low responsiveness to blockade seen in T_FH_, we examined expression of the other ICs TIGIT and CD200, which are also frequently expressed on T_FH_ cells (34). Similar to PD-1, expression of these IC was higher on AIM+ HIV-specific CD4+ T cells of CP than EC (Fig 5a-d), and correlated positively with viremia (Fig S5a), demonstrating an association between antigen burden and their accumulation, as previously shown for PD-1 (16) and TIGIT (35). Single-cell expression of TIGIT and CD200 was directly correlated with that of PD-1 (Fig 5ef), as well as with each other (Fig S5b). Almost half of AIM+ HIV-specific CD4+ T cells of CP co-expressed all three IC, whereas only a small fraction was triple-positive in EC (Fig 5g), in line with IC co-upregulation in conditions of elevated CD4+ T cell dysfunction (10). These IC varied according to subsets of HIV-specific CD4+ T cells (Fig S5cd). TIGIT was high on CXCR5+ cells and low on CCR6+ cells in both cohorts, and high on the CXCR3+ of CP only (Fig S5e). CD200 expression followed very similar patterns (Fig S5f). Consistently, IC expression differed between HIV-specific cytokine mRNA+ CD4+ T cells of CP, with high expression on *IL21, CXCL13, IFNG* and *IL2* mRNA+ cells, and low IC levels on *GZMB, IL17F* and *IL22* mRNA+ cells (Fig 5hi). Notably, CD200 was undetectable on mucosal-related cytokine mRNA+ cells. These patterns were conserved in EC (Fig S5ef), with once again the exception of *IFNG* mRNA+ cells, on which TIGIT and CD200 levels were low. Thus, TIGIT and CD200 are highly expressed on HIV-specific CD4+ T cells producing IL-2 or T_FH_-associated cytokines even in the absence of high viral loads, yet IC accumulate on other cytokine+ cells only in the setting of dysfunction, in particular for functions reduced in CP compared to EC.

**Figure 5.**
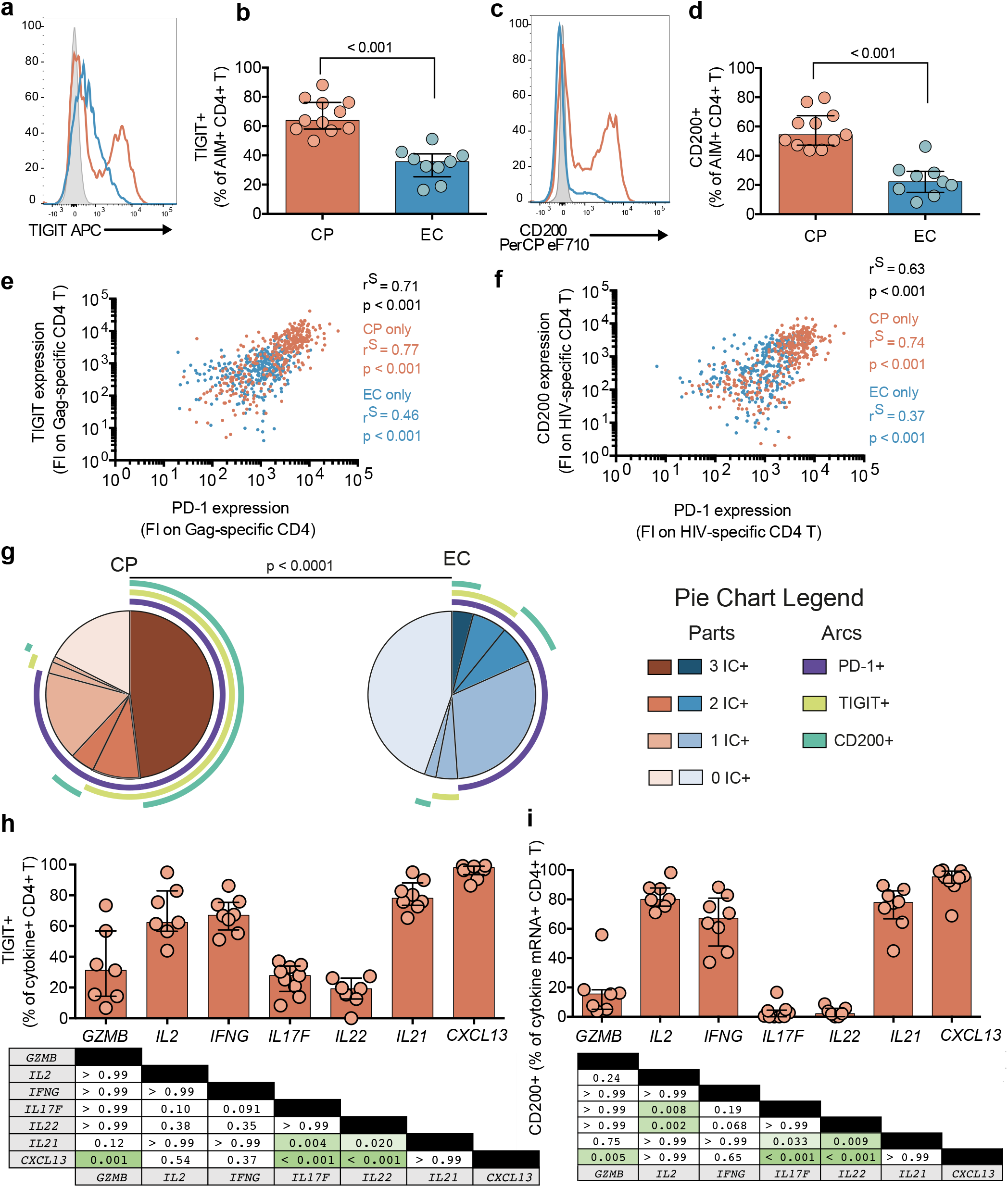
Differential expression of ICs among functionally-distinct subsets of HIV-specific CD4+ T cells. Representative histogram overlays of **a**) TIGIT or **c**) CD200 expression on AIM+ Gag-specific CD4+ T cells from a CP (red) or an EC (blue). Grey shaded outline represents PD-1 FMO. Fraction of AIM+ Gag-specific CD4+ T cells expressing **b**) TIGIT or **d**) CD200 of either cohort. Correlation between single-cell expression of PD-1 and **e**) TIGIT or **f**) CD200 on AIM+ Gag-specific CD4+ T cells from 4 CP and 4 EC (100 cells per subject). **g**) Co-expression patterns between the ICs PD-1, TIGIT and CD200 on AIM+ HIV-specific CD4+ T cells from both cohorts. Shades of pie parts represent number of ICs; arcs represent IC expressed in pie part. Cumulative data of **h**) TIGIT and **i**) CD200 expression on cytokine mRNA+ Gag-specific CD4+ T cells from CP. Statistics appear in tables below, with p values < 0.05 highlighted in green. bd) n = 13 CP and 8 EC, only positive responses (at least 2 fold greater than unstimulated) were considered; g) n = 9 CP, only positive responses were considered. Columns and pie chart fractions correspond to median values, with interquartile range for columns. bd) Mann-Whitney test; ef) Spearman correlation; g) permutation test with 10 000 permutations; hi) Friedman test with Dunn’s post-test. FI = Fluorescence Intensity.

Combined ICB strategies targeting different molecules can be more potent than single blockade (10, 11). We examined the impact of two clinical-grade ICB antibodies developed for immunotherapy, the anti-PD-L1 antibody BMS-936559 and the anti-TIGIT antibody BMS-g86207-Ab (Bristol-Myers Squibb) using d-ICS, allowing us to multiplex the four cytokines IFNγ, IL-2, IL-21 and CXCL13 (Fig S5i). Single TIGIT blockade did not increase cytokine+ responses for any of the functions studied (Fig S5j). Responses were heterogeneous within the CP cohort: depending on the participant, we observed limited response to any blockade strategy (Fig S5h, left), detectable responses in the co-blockade condition only (Fig S5h, middle) or modest to no benefit of co-blockade compared to single PD-1 blockade (Fig S5h, right). Our data suggests co-blockade strategies may generate responses in a larger fraction of individuals than single-blockade, although some subjects may remain unresponsive.

Because of the differential response of IFNγ and CXCL13 to blockade, we compared its impact between monofunctional cells and the population co-expressing CXCL13 and IFNγ, which represented roughly 20% of the overall cytokine-producing populations (Fig S5i). The frequency of IFNγ single-positive and or double-positive CD4+ T cells increased upon PD-L1 blockade, whereas CXCL13 single-positive cells remained refractory (Fig S5jk). The double-positive population are also increased upon blockade, highlighting that CXCL13 transcription can be susceptible to ICB. The pattern was the same with co-blockade, at a slightly greater magnitude. These observations highlight the accumulation of ICs TIGIT and CD200 on subsets other than TFH in the context of dysfunction. However, high co-expression of TIGIT and PD-1 does not result in enhanced response to TIGIT and PD-L1 co-blockade compared to single PD-L1 blockade in most individuals.

### ART-induced viral suppression differentially affects HIV-specific CD4+ T cell response to ICB

As ICB in HIV infection is predominantly being evaluated in ART-suppressed individuals, we compared responses in longitudinal samples obtained before and after ART (Fig 6a, Fig S6a). ART initiation led to a consistent decrease of the preferentially T_FH_-associated functions IL-21 and CXCL13 in HIV-specific CD4+ T cells (Fig 6b). With the exception of the frequency of *IL2* mRNA+ CD4+ T cells, which remained constant between both time points, the effect of ART was heterogeneous and subject-dependent, with some subjects experiencing an increase or a decrease, and others still maintaining stable responses (Fig 6b). Post-ART, IC expression decreased on all cytokine mRNA+ CD4+ T cells, except for the *IL2*+, on which IC expression was maintained (Fig S6b-d). These results highlight a correction of the high IC expression and T_FH_-like skewing acquired in viremia towards a profile more similar to that observed in EC, while other functions are inconsistently recovered.

**Figure 6.**
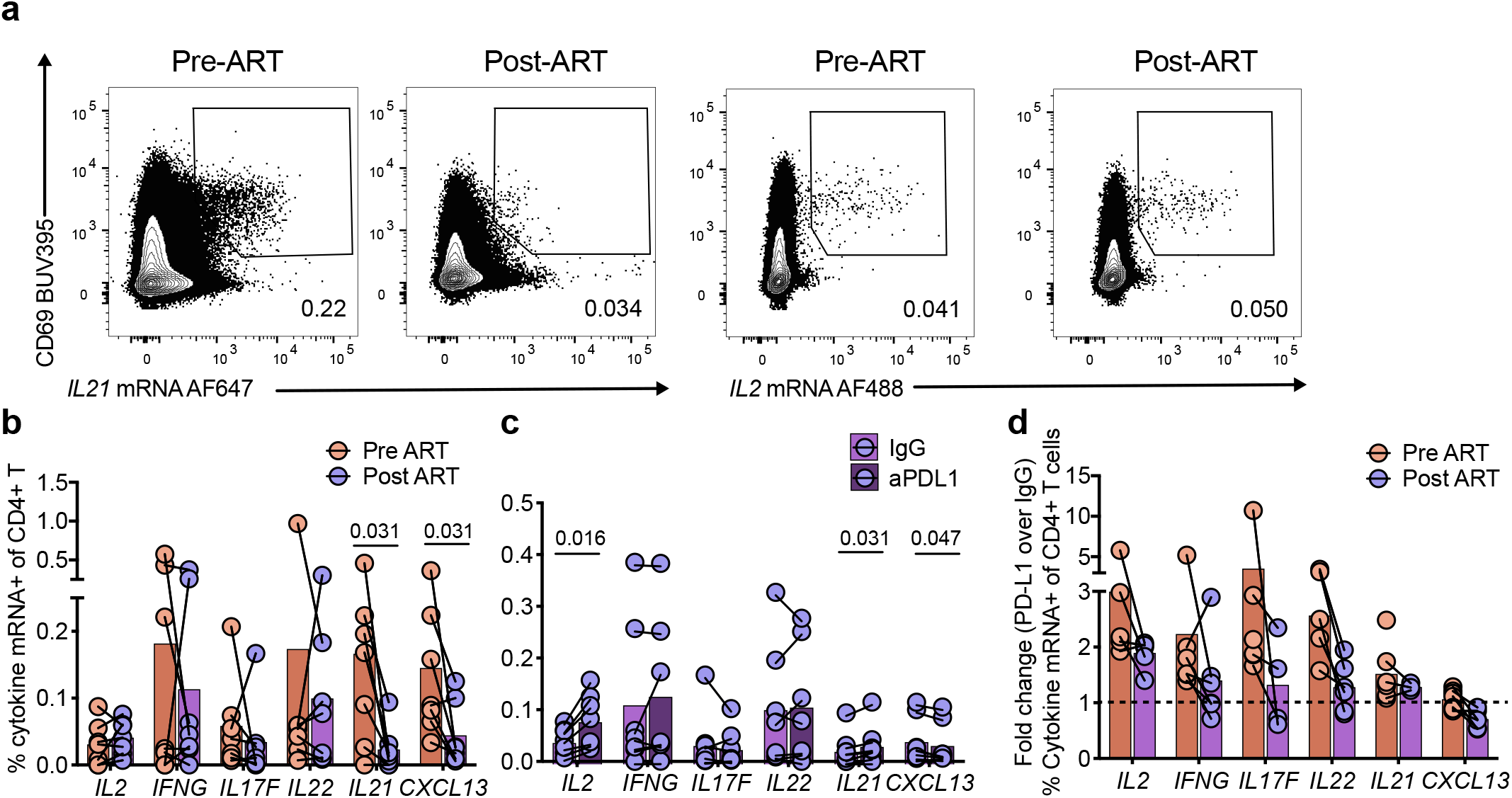
Viral suppression on ART differentially affects responsiveness of effector functions to PD-L1 blockade. **a)** Representative examples and **b)** summary data of net frequency of Gag-specific cytokine mRNA+ CD4+ T cells from matched subjects prior to ART (orange) and after ART (purple) following a 9hr in vitro stimulation with Gag peptide pool. **c)** Summary net frequencies of Gag-specific cytokine mRNA+ CD4+ T cells stimulated with PD-L1 blocking antibody or isotypic control (IgG) among ART-treated individuals. **d**) Comparison of the fold changes upon PD-L1 blockade between longitudinal samples pre- (orange) and post-ART (purple). N = 7 longitudinal samples. Bars represent medians. bcd) Wilcoxon test.

The increase of *IL2* mRNA+ CD4+ T cells upon PD-L1 blockade was also seen following ART treatment (Fig 6c), and of similar magnitude to that observed in both CP (Fig 6d) and EC (Fig 4d). However, the increases for *IFNG* and mucosal-related cytokine expression were generally less pronounced after ART initiation, albeit with inter-subject variability (Fig 6cd). No effect on the frequency of T_FH_-cytokine+ CD4+ T cells was observed upon blockade during ART.

These data suggest that increased IL-2 production is a maintained benefit of PD-L1 blockade, while the increase in *IFNG* and mucosal-related cytokines upon ICB is subdued once ART is initiated.

## Discussion

Immune checkpoints inhibit T cell activation through multiple mechanisms (36, 37). Although recent findings have partially elucidated the molecular features behind these effects (5, 37), an understudied topic remains if and how these immune checkpoints operate differently among the heterogeneous lineages of CD4+ T cells. Using high-parameter flow cytometry combining protein and FISH mRNA staining, we assessed HIV-specific CD4+ T cells of an array of T helper phenotypes and functions otherwise difficult to measure. We focused on a palette of T_FH_, T_H_1 and T_H_17/T_H_22-associated traits. These phenotypes, as identified by canonical chemokine receptors and transcription factors or by production of effector molecules, presented a hierarchy of relative expression levels of TOX, PD-1, TIGIT and CD200. This differential expression was present at all HIV disease stages, although the magnitude of expression was associated with viral load. Responsiveness to PD-L1 blockade varied according to a defined function of CD4+ T cells rather than their levels of IC expression. PD-L1 blockade had more limited effects in individuals with spontaneous or therapeutic control of viral replication than in people with high antigen load. These data highlight a previously unappreciated heterogeneity of responsiveness to ICB among HIV-specific CD4+ T cells.

TOX expression was greater in HIV-specific CD4+ T cells of CP compared to EC, in line with their greater state of dysfunction linked with ongoing antigen stimulation in CP, and greater functionality in EC(4). TOX expression is linked to repeated TCR stimulation (7, 38), a central driver of T cell exhaustion, and was strongly associated to PD-1 levels in the presence of viremia. HIV-specific CD4+ T cells from a same subject expressed different amounts of IC depending on their polarization, consistent across function-dependent (ICS) and function-agnostic (AIM) methods of identification. High expression or co-expression of IC did not prevent effector functions, as observed with IC-high CXCL13+, IL-21+ and IL-2+ cells. IL-2 markedly increased with PD-L1 blockade, consistent with an inhibitory effect by PD-1, while no effect was observed for IL-21 and CXCL13. The hierarchy of IC expression between polarizations of HIV-specific CD4+ T cells suggest IC may not equally regulate the respective’ CD4+ T cell function. As shown in a TCR transfection model system of primary human PBMCs, some T cell functions are more resistant to PD-1-mediated inhibition than others (39), while a recent study using mouse and human T cell lines demonstrated different sensitivities of gene expression to PD-1 inhibition (40). IC may not be inhibitory in all instances: T_FH_ express lower amounts of IL-21 and IL-4 following PD-1 ablation in mice (41); TIGIT, although inhibitory when expressed on CD8+ or T_H_1 T cells (42, 43), is associated with strong B cell help and cytokine expression in T_FH_ (44); CD200 is associated with lack of pro-inflammatory cytokines, yet high IL-4 production in CD4+ T cells (45). Although these reports often find IC not inhibiting T_FH_-related functions, our observation that CXCL13+IFNγ+ cells increased in frequency upon ICB indicates this T_FH_ function can be negatively modulated by PD-1. Co-expression of CXCL13+ cells with a T_H_1-associated cytokine may correspond to a transition into a more plastic cell state which is responsive to ICB, while the absence of response in CXCL13 single-positive cells suggest that the cell-intrinsic state is associated with response to blockade, rather than single cytokine pathways or IC expression.

Response to PD-L1 blockade was stronger in both breadth and magnitude for the dysfunctional HIV-specific CD4+ T cells of the CP compared to the EC and ART cohorts, suggesting that antigen presence sensitizes antigen-specific CD4+ T cells to ICB. Co-blockade with a TIGIT-blocking antibody further enhanced the effect of PD-L1 blockade only in some patients, consistent with the reported varying sensitivity to co-blockade among subjects (12, 35, 46), and highlighting the central inhibitory role of PD-1. Of note, HIV-specific CD4+T cells may respond directly to ICB by blockade of autologous PD-1 molecules, as we have shown with live-sorted CD4 T cells subsets and add back co-culture experiments(14), or indirectly by paracrine mechanisms, like the feed forward loop of soluble factors between T cells and antigen-presenting cells (47). While their respective contributions would be extremely challenging to delineate on primary human T cells, the critical observation remains that different HIV-specific CD4 T cell subsets have a differential ability to respond to PD-1 blockade.

CD4+ T cells expressing mucosal cytokines responded well to PD-L1 blockade, despite the low levels of PD-1 expression on these cells overall. This suggests the replenished gut in the chronic SIV model may be linked to responsiveness of T_H_17/T_H_22 cells (25, 26), the primary CD4+ T cell population of that anatomical site. In this scenario, bacteria-specific T_H_17 may also have responded to ICB (48). Taken together, our data suggests that polarizations nudging towards T_H_1 and T_H_17 may undergo a more direct inhibition by PD-1, explaining their strong responsiveness to ICB. This can procure benefits such as direct antiviral control and restored gut integrity, even under conditions of persistent antigen.

As rapid initiation of ART is now the standard of care upon HIV diagnosis, it is crucial to know whether the response to PD-L1 blockade changes once viremia is therapeutically suppressed. IL-2 response reached the same magnitude as observed at the pre-ART time point, comparable to that seen in EC, in line with the direct inhibitory role PD-1 plays in regulating IL-2 (20). Conversely, responses of CD4+ T cells expressing IFNγ and mucosal-related cytokines to ICB decreased in magnitude once ART was initiated. The general lowering of reactivity to ICB in contexts of controlled viremia strongly support a role of ongoing antigen presence in sensitizing these cells to this type of treatment. In addition, ART may block *de novo* virus-specific CD4+ T cells, which may be more responsive to ICB (49). These observations highlight the important role timing may play to maximize benefits of ICB in the context of HIV.

Although more comprehensive than previous studies looking at HIV-specific CD4+ T cells, our observational study focused on a selected set of T helper functions. Furthermore, while we utilized *in vitro* blockade of PBMCs to interrogate the response of multiple CD4+ T cell subsets to PD-L1 blockade, most HIV-specific CD4+ T cells reside in tissues, where IC can play distinct roles, and cannot address the actively-researched *in vivo* role of ICB(50). Addressing such questions would require tissue samples or animal models. Indeed, a clinical safety trial of anti-PD-L1 in ART-treated revealed 2 of 6 HIV-infected individuals had increased HIV-specific CD8+ T cell responses in blood (51). In the non-human primate model (NHP) of Simian Immunodeficiency Virus (SIV) infection, PD-1 blockade did significantly decreased viral load in chronically-infected NHPs (24). In ART-treated NHPs, PD-1 blockade and PD-1/CTLA dual blockade resulted in a 5-day delay in viral rebound following ART cessation, but was not sufficient to achieve viral control (52), suggesting ICB may have to be combined with additional strategies. Indeed, combination of ART, an anti-SIV-boosting vaccine and PD-1 blockade was recently shown to suppress rebound after ART interruption in rhesus macaques (53). These approaches should be considered with precaution, as restoring function of virus-specific T cells can cause depletion of lymphoid organs harboring infected cells, impeding the generation of new immune responses (54).

In summary, we highlight an intrinsic heterogeneity in IC expression among different polarizations of HIV-specific CD4+ T cells, revealing a disconnect between classical notions of IC and their relevance among CD4+ T cells lineages. This data also shows that functional lineages of HIV-specific CD4+ T cells have different capacities to respond to IC blockade, with CD4+ T cells expressing mucosal-protective and antiviral-associated cytokines responding well, whereas T_FH_-associated cytokines responded poorly. These results emphasize the importance of considering CD4+ T cell differentiation in studies of IC blockade in the context of T cell dysfunction, and may have implications for CD4+ T cell help in other infectious and non-infectious chronic human diseases.

## Acknowledgments

We thank Josée Girouard, the clinical staff at the McGill University Health Centre in Montreal and all study participants for their invaluable role in this project. We also thank Ms. Alina Dyachenko for help with statistical analyses.

## Contributors

<u>Conceptualization:</u> EBR, DEK, AM, MD

<u>Data Curation</u>: EBR, AM, JN

<u>Formal analysis</u>: EBR, AM, OT, DEK

<u>Funding acquisition</u>: DEK, MD

<u>Investigation/experiments:</u> EBR, AM, NB, GOD, RC

<u>Methodology:</u> EBR, AM, JN, AEB

<u>Resources</u>: GJF, JPR, CT

<u>Software:</u> OT

<u>Visualization:</u> EBR, OT

<u>Patient recruitment and cohort administration:</u> NB, CT, JPR

<u>Supervision:</u> DEK, AM, MD, AEB

<u>Writing – original draft:</u> EBR, AEB, DEK

<u>Writing – review and editing</u>: all authors

