## Supplemental materials for "Immune checkpoint expression on HIV-specific CD4+ T cells and response to their blockade are dependent on lineage and function"

### Supplementary Materials

Supplemental Figure 1. Transcriptomic analysis of exhaustion-related transcription factors in HIV-specific CD4+ T cells.
Supplemental Figure 2. PD-1 expression on subsets of AIM+ HIV-specific CD4+ T cells.
Supplemental Figure 3. Transcription of multiple Thelper cytokines is detected by RNA-Flow-FISH following HIV-specific stimulation
Supplemental Figure 4. Protein data corroborates cytokine mRNA responses of HIV-specific CD4+ T cells.
Supplemental Figure 5. Despite high ICs' TIGIT and CD200 expression among polarizations of HIV-specific CD4+T cells, co-blockade of PD-1 and TIGIT does not increase responsiveness to ICB.
Supplemental Figure 6. ART initiation causes a decrease of IC upon HIV-specific cytokine mRNA+ CD4+ T cells.
Supplementary table 1: Subject characteristics
Supplementary Table 2: Flow cytometry panel used in AIM assay
Supplementary Table 3: Variants of the Flow cytometry panels used in intra-nuclear transcription factor staining
Supplementary Table 4: Flow cytometry panel used in mRNA-Flow-FISH assay
Supplementary Table 5: Probe panel combination used in mRNA-Flow-FISH assay
Supplementary Table 6: Flow cytometry panel used in delayed intracellular cytokine staining (d-ICS)

**Supplemental Figure 1. Transcriptomic analysis of exhaustion-related transcription factors in HIV-specific CD4+ T cells. a**) Representative flow cytometry plots of the full gating strategy for the AIM panel on HIV Gag-stimulated cells from a CP. Gating strategy includes CD3-low CD69+ as CD3 expression can be downregulated following TCR stimulation. **b**) Representative example of PD-1 expression on AIM+ Gag-specific CD4+ T cells. Black line is FMO. **c**) Relative mRNA expression of exhaustion-related TF among sorted Gag-specific CD4+ T cells of CP (red) or EC (blue), as captured by high-throughput RT-PCR (Fluidigm® - for details, please see (4). n = 12 CP and 12 EC. **d**) Histogram overlay of flow cytometry data showing the expression of TF in AIM+ Gag-specific CD4+ T cells of two CP of varying viral loads (red, orange) and one EC (blue). Dotted line is expression in unstimulated naïve CD4+ T cells. E) Expression of TOX among total unstimulated CD4+ T cells per cohort. c) n = 12 CP and 12 EC. e) n = 8 CP and 8 EC. ce) Stats = Mann-Whitney test.

**Supplemental Figure 2. PD-1 expression on subsets of AIM+ HIV-specific CD4+ T cells**. **a**) Summary of expression of the chemokine receptors CXCR3, CCR6 and CXCR5 on AIM+ Gag-specific CD4+ T cells from CP (orange) or EC (blue). **b**) Representative flow cytometry plots depicting gating strategy to identify TH17 (CCR6+RORgt+CXCR3-), TH1/TH17 (CCR6+RORgt+CXCR3+), TH1 (CXCR3+CCR6-Tbet+Eomes+) or TFH (CXCR5+BCL-6+), either in total CD4+ (top row) or AIM+ Gag-specific CD4+ T (bottom row). **c**) Cumulative data of the frequency of TF-identified subsets among AIM+ Gag-specific CD4+ T cells. a) n= 13 CP and 9 EC; c) n = 8 CP and 8 EC. Columns correspond to median values with interquartile range. ac) Mann-Whitney test.

**Supplemental Figure 3. Transcription of multiple Thelper cytokines is detected by RNA-Flow-FISH following HIV peptide stimulation**. **a**) Representative flow cytometry plots of the full gating strategy for the RNA-Flow-FISH panels (see methods for details). **b**) Representative examples of cytokine mRNA+ Gag-specific CD4+ T cells as detected by RNA-Flow-FISH following a 12hr Gag stimulation for a CP. Gates set on unstimulated (Unstim) conditions. **c**) Cumulative data for detection of *IL4* mRNA and *IL10* mRNA production following Gag stimulation in both cohorts. Negative responses (less than 2 fold over unstimulated condition) are identified by grey symbols. **d**) Response of all cytokine mRNA combinations per panel to Gag-stimulation in CP compared to EC. **e**) Euler diagrams of chemokine receptor expression among HIV-specific CD4+ T cells identified by production of *GZMB*, *IL2*, *IFNG*, *IL17F*, *IL22*, *IL21* or *CXCL13* mRNA. Values represent median frequencies of subsets among cytokine mRNA+ Gag-specific CD4+ T cells from 9 CP (red hues) or 8 EC (blue hues). n = 9 CP and 8 EC. Columns correspond to median values with interquartile range. Mann-Whitney, n= 9 CP and 8 EC.

**Supplemental Figure 4. Protein data corroborates cytokine mRNA responses of HIV-specific CD4+ T cells. a**) Representative flow cytometry plots showing detection of protein cytokine production by delayed ICS following stimulation with a HIV Gag peptide pool in a CP. **b**) Correlation between the net frequency of Gag-specific CD4+ T cells, as detected by cytokine mRNA+ cells by RNA FlowFISH (x axis) or by protein cytokine assessed by delayed ICS (y axis) for IL-2 (left), IFNγ (midde) or CXCL13 (right) in CP. n= 8 CP (only donors with Gag-specific responses at least 2-fold over unstimulated are shown). Statistical comparison: Spearman correlation. **d**) Frequency of protein cytokine+ cells detected by delayed ICS upon Gag stimulation, either with isotype control (IgG) or with PD-L1 blocking antibody (aPD-L1). Response of all cytokine mRNA combinations to PD-L1 blockade, upon Gag-stimulation among EC, for **e**) antiviral panel, **f**) mucosal panel, or **g**) TFH panel. n = 8 EC. Statistical comparison: Wilcoxon test. Fold change in the net frequencies of cytokine mRNA+ Gag-specific CD4+ T cells detected upon PD-L1 blockade compared to isotypic control for both cohorts for **h**) antiviral panel, **i**) mucosal panel, or **j**) TFH panel. CP in orange and EC in blue. n= 9 CP and 8 EC (only donors with Gag-specific responses at least 2-fold over unstimulated are shown). Statistical comparison: Mann-Whitney test. Each donor within a cohort has been separately color coded. Columns correspond to median values with interquartile range.n = 10 CP. Stats : Wilcoxon test.

**Supplemental Figure 5. Despite high ICs' TIGIT and CD200 expression among polarizations of HIV-specific CD4+T cells, co-blockade of PD-1 and TIGIT does not increase responsiveness to ICB**. **a)** Correlations between viral load of CP and their IC expression (PD-1 gMFI on left, % TIGIT+ in middle and % CD200+ on right) Gag-specific AIM+ CD4+ T cells. **b**) Correlation between single-cell expression of TIGIT and CD200 on AIM+ HIV-specific CD4+ T cells from 4 CP and 4 EC (100 cells per subject). Frequency of **c**) TIGIT+ or **d**) CD200+ among the three polarizations of the AIM+ Gag-specific CD4+ T cells defined by CXCR3, CCR6 and CXCR5 expression in both cohorts – CP (red) and EC (blue). Grey circles represent negative responses (less than 2 fold over unstimulated condition). Cumulative data of **e**) TIGIT and **f**) CD200 expression on cytokine mRNA+ Gag-specific CD4+ T cells from EC cohort. Statistics appear in tables below, with p values $< 0.05$ highlighted in green. **g**) Response (fold change compared to IgG) to all blocking strategies as detected by delayed ICS in CP. **h**) Responses of individuals' responses (fold change compared to IgG) with no benefit of dual blockade over single PD-L1 blockade (left), further enhancement of cytokine production by dual blockade over single PD-L1 blockade (middle), or no (or modest) effect of either blockade strategy (right). **i**) Venn representation of IFNγ and CXCL13 median co-expression in the CP cohort. **j**) Net frequencies of IFNγ single-positive (SP), CXCL13 SP or IFNγ/CXCL13 double-positive Gag-specific CD4+ T cells following stimulation in the presence of IgG or aPD-L1. Statistical comparison by Wilcoxon test. **k**) Fold change of blocking conditions (compared to IgG) for IFNγ SP, CXCL13 SP or DP. bh) Statistical comparison by 2-way ANOVA with Tukey's multiple comparison test. a, c-f) n= 13 CP, 9 EC. g-k) N = 10 CP. Bars represent medians with interquartile range. de) N = 13 CP and 8 EC; fg) n = 8 EC, where only positive responses are considered. Friedman's test with Dunn's correction. abc) Spearman R correlation. Viral load was Log10 transformed.

**Supplemental Figure 6. ART initiation causes a decrease of IC upon HIV-specific cytokine mRNA+ CD4+ T cells. a**) gMFI of cytokine mRNA+ in Gag-specific cytokine mRNA+ CD4+ T cells from matched subjects prior to ART (ored) and after ART (purple) following a 9hr *in vitro* stimulation with Gag peptide pool. Comparison of IC expression among Gag-specific cytokine mRNA+ CD4+ T from matched donors: **b**) PD-1 gMFI, **c**) frequency of TIGIT+, or **d**) frequency of CD200+. Only positive responses (at least 2 fold over unstimulated condition) were considered. bcd) Wilcoxon test, n = 7 longitudinal samples. gMFI = geometric mean fluorescence intensity.

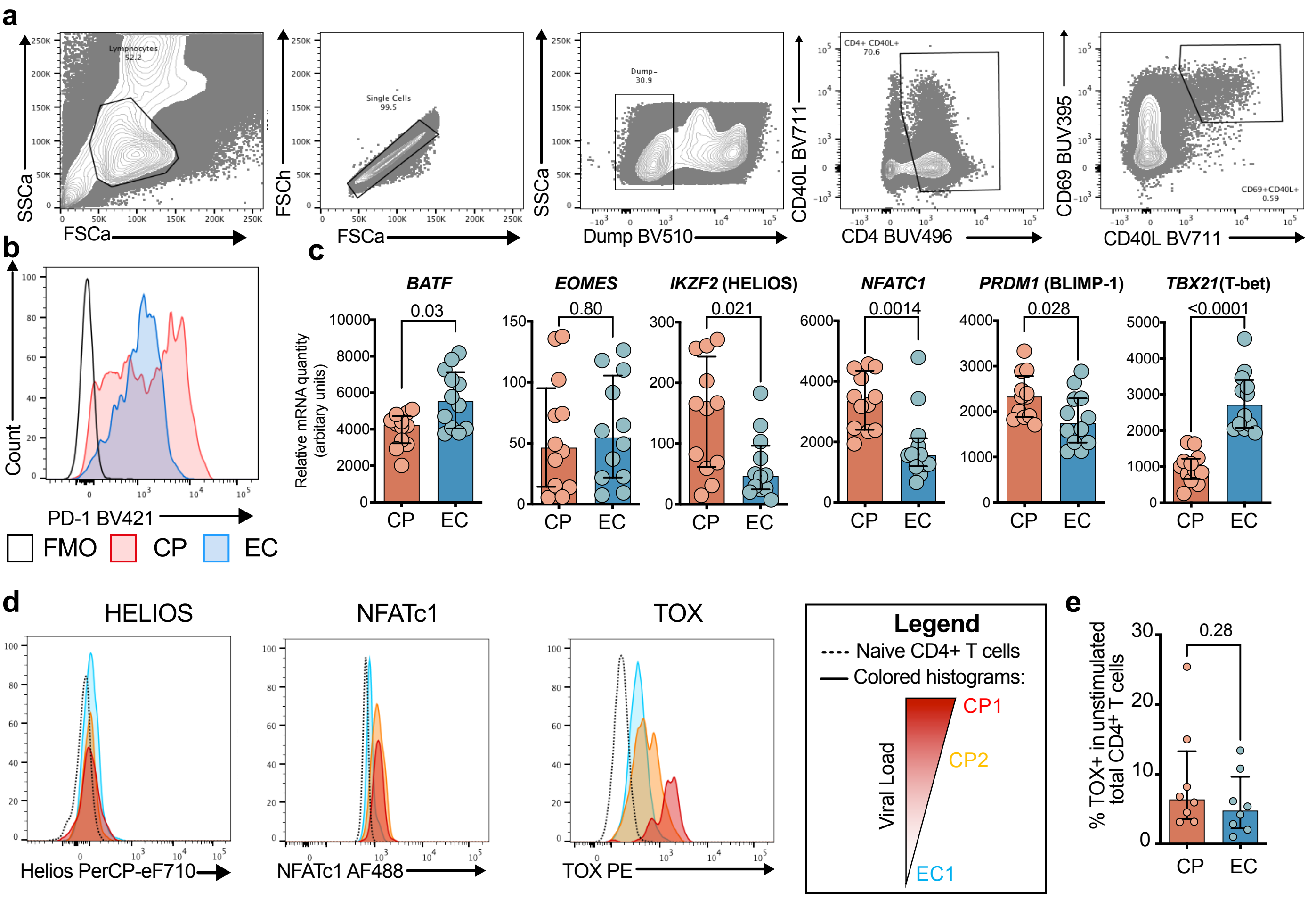
a
Lymphocytes 52.2
Single Cells 99.5
Dump- 30.9
CD4+ CD40L+ 70.6
CD69+CD40L+ 0.59
SSCa
FSCa
FSCh
Dump BV510
CD40L BV711
CD4 BUV496
CD69 BUV395
b
Count
PD-1 BV421
FMO
CP
EC
c
Relative mRNA quantity (arbitary units)
BATF
0.03
EOMES
0.80
IKZF2 (HELIOS)
0.021
NFATC1
0.0014
PRDM1 (BLIMP-1)
0.028
TBX21(T-bet)
<0.0001
CP
EC
d
HELIOS
NFATc1
TOX
Helios PerCP-eF710
NFATc1 AF488
TOX PE
Legend
Naive CD4+ T cells
Colored histograms:
Viral Load
CP1
CP2
EC1
e
% TOX+ in unstimulated total CD4+ T cells
0.28
CP
EC

**S1**

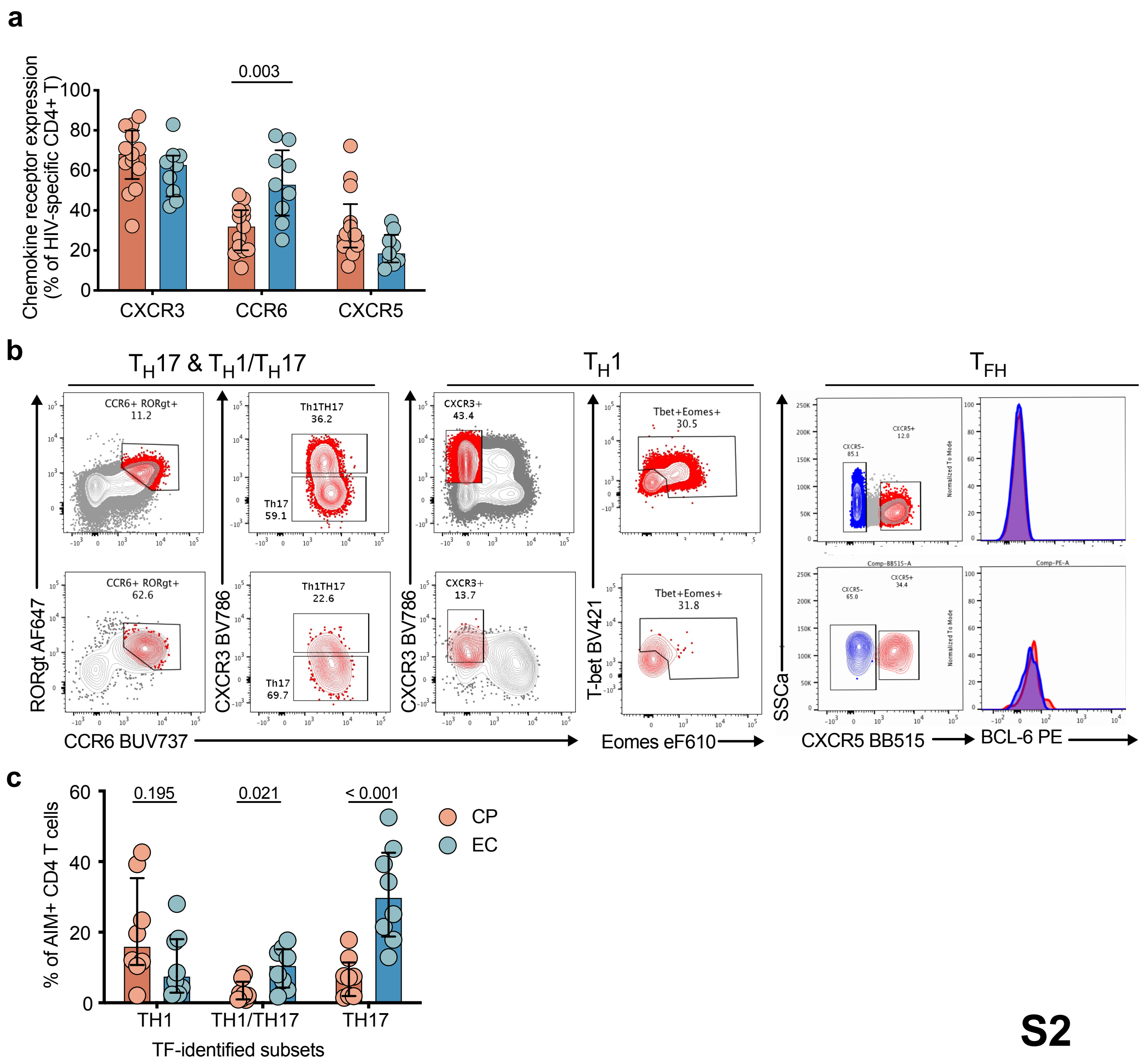
a
Chemokine receptor expression (% of HIV-specific CD4+ T)
0.003
CXCR3
CCR6
CXCR5
b
TH17 & TH1/TH17
TH1
TFH
CCR6+ RORgt+ 11.2
Th1TH17 36.2
Th17 59.1
CXCR3+ 43.4
Tbet+Eomes+ 30.5
CCR6+ RORgt+ 62.6
Th1TH17 22.6
Th17 69.7
CXCR3+ 13.7
Tbet+Eomes+ 31.8
RORgt AF647
CXCR3 BV786
T-bet BV421
SSCa
CCR6 BUV737
Eomes eF610
CXCR5 BB515
BCL-6 PE
c
% of AIM+ CD4 T cells
0.195
0.021
< 0.001
CP
EC
TH1
TH1/TH17
TH17
TF-identified subsets

**S2**

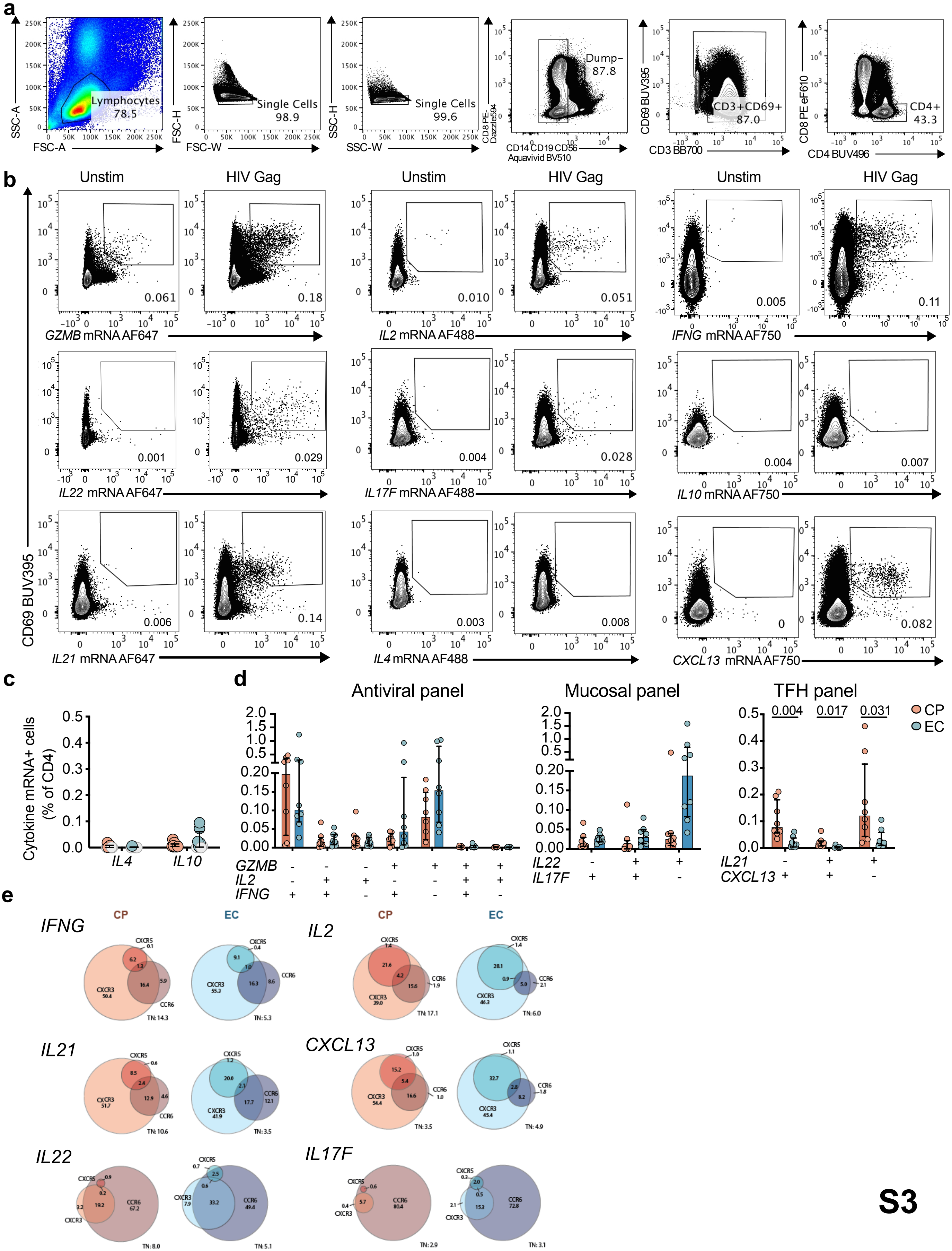

a
Lymphocytes 78.5
Single Cells 98.9
Single Cells 99.6
Dump- 87.8
CD3+CD69+ 87.0
CD4+ 43.3
SSC-A
FSC-A
FSC-H
FSC-W
SSC-H
SSC-W
CD8 PE-Dazzle594
CD14 CD19 CD56 Aquavivid BV510
CD69 BUV395
CD3 BB700
CD8 PE eF610
CD4 BUV496
b
Unstim
HIV Gag
CD69 BUV395
GZMB mRNA AF647
IL2 mRNA AF488
IFNG mRNA AF750
IL22 mRNA AF647
IL17F mRNA AF488
IL10 mRNA AF750
IL21 mRNA AF647
IL4 mRNA AF488
CXCL13 mRNA AF750
c
Cytokine mRNA+ cells (% of CD4)
IL4
IL10
d
Antiviral panel
Mucosal panel
TFH panel
CP
EC
0.004
0.017
0.031
GZMB
IL2
IFNG
IL22
IL17F
IL21
CXCL13
e
IFNG
IL2
IL21
CXCL13
IL22
IL17F
CXCR5
CXCR3
CCR6
S3

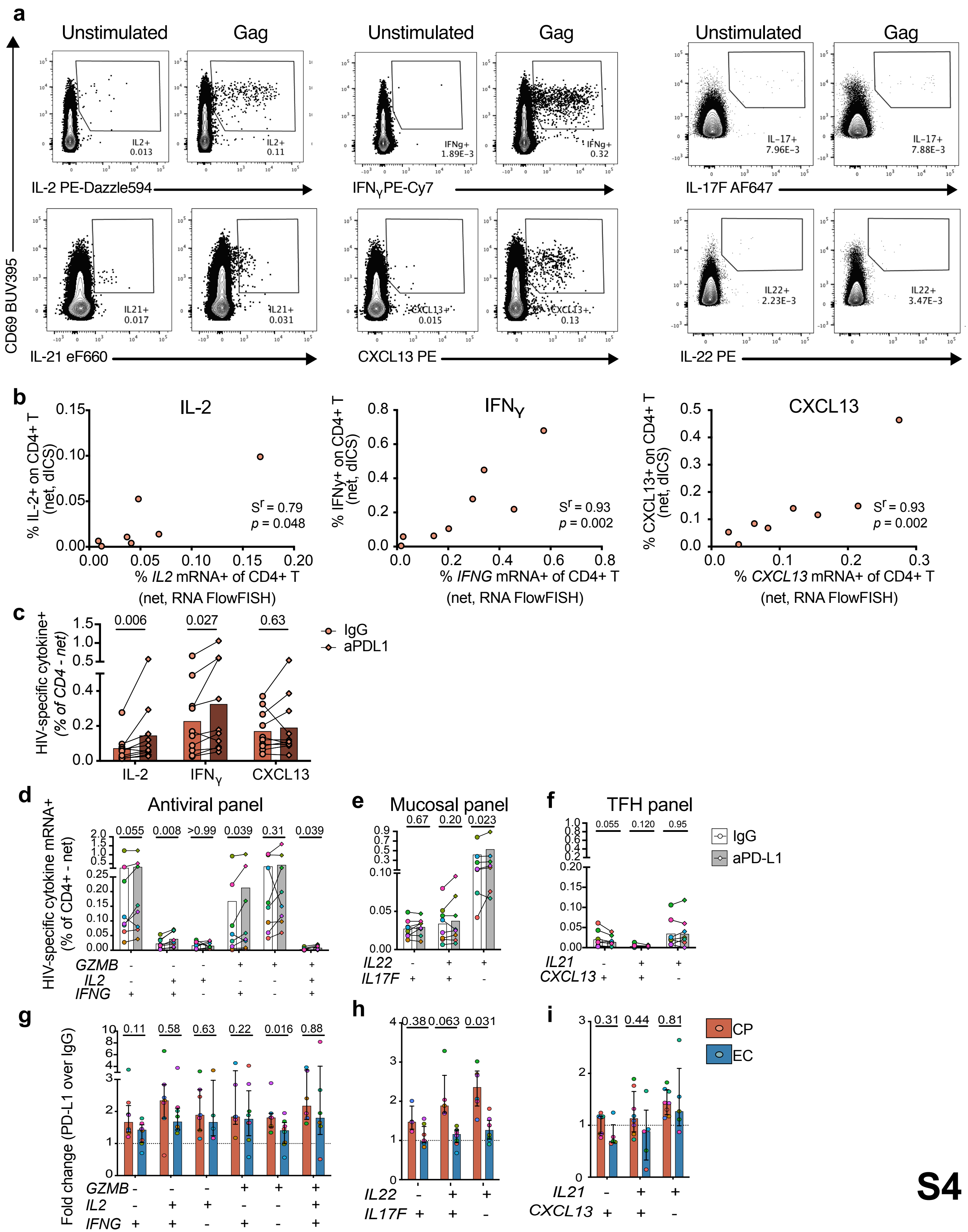

a
Unstimulated
Gag
IL2+ 0.013
IL2+ 0.11
IL-2 PE-Dazzle594
IFNg+ 1.89E-3
IFNg+ 0.32
IFNγPE-Cy7
IL-17+ 7.96E-3
IL-17+ 7.88E-3
IL-17F AF647
CD69 BUV395
IL21+ 0.017
IL21+ 0.031
IL-21 eF660
CXCL13+ 0.015
CXCL13+ 0.13
CXCL13 PE
IL22+ 2.23E-3
IL22+ 3.47E-3
IL-22 PE
b
IL-2
% IL-2+ on CD4+ T (net, dICS)
% IL2 mRNA+ of CD4+ T (net, RNA FlowFISH)
Sr = 0.79 p = 0.048
IFNγ
% IFNγ+ on CD4+ T (net, dICS)
% IFNG mRNA+ of CD4+ T (net, RNA FlowFISH)
Sr = 0.93 p = 0.002
CXCL13
% CXCL13+ on CD4+ T (net, dICS)
% CXCL13 mRNA+ of CD4+ T (net, RNA FlowFISH)
Sr = 0.93 p = 0.002
c
HIV-specific cytokine+ (% of CD4 - net)
0.006
0.027
0.63
IgG
aPDL1
IL-2
IFNγ
CXCL13
d
Antiviral panel
HIV-specific cytokine mRNA+ (% of CD4+ - net)
0.055
0.008
>0.99
0.039
0.31
0.039
GZMB
IL2
IFNG
e
Mucosal panel
0.67
0.20
0.023
IL22
IL17F
f
TFH panel
0.055
0.120
0.95
IL21
CXCL13
IgG
aPD-L1
g
Fold change (PD-L1 over IgG)
0.11
0.58
0.63
0.22
0.016
0.88
GZMB
IL2
IFNG
h
0.38
0.063
0.031
IL22
IL17F
i
0.31
0.44
0.81
IL21
CXCL13
CP
EC


S4

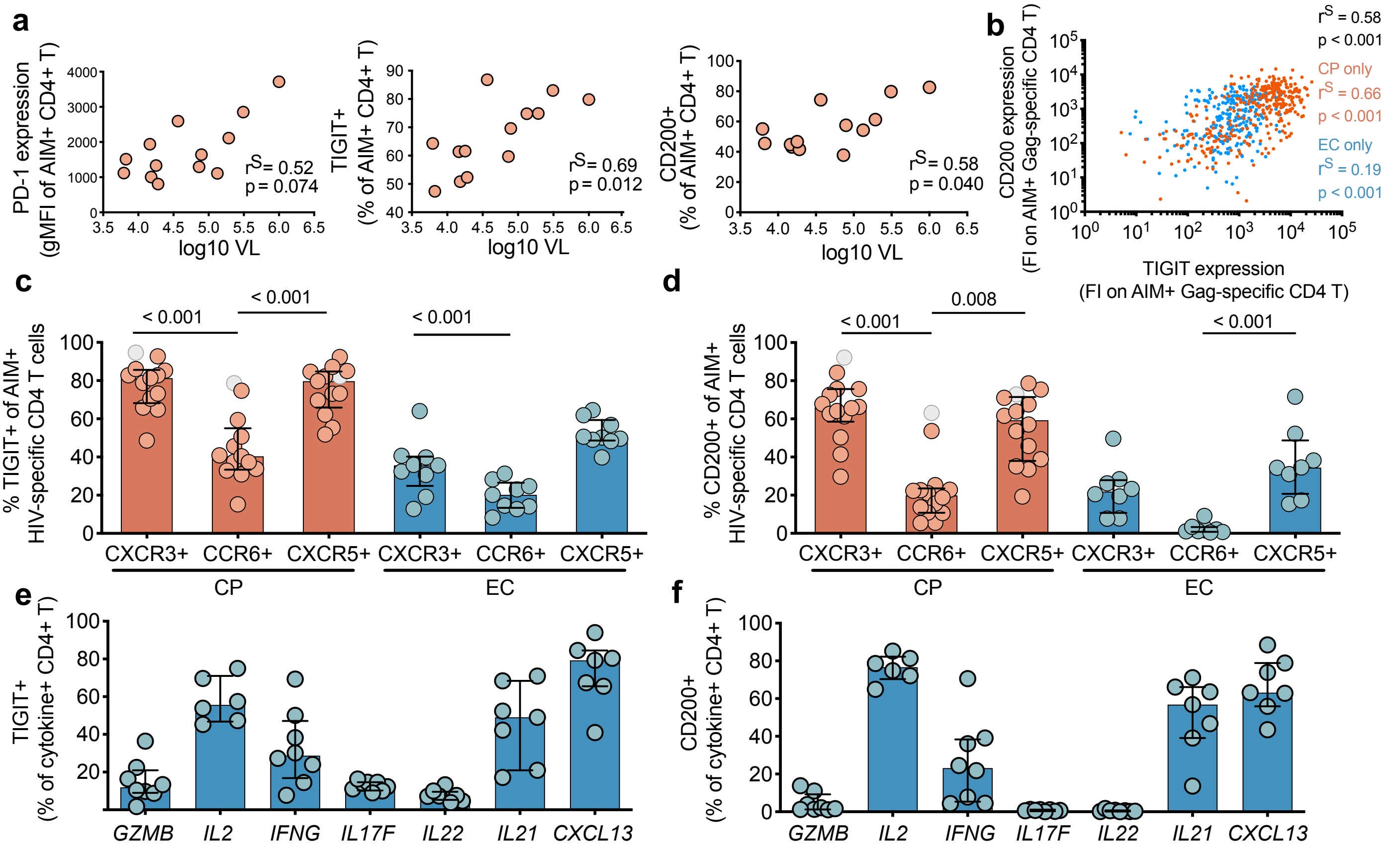


| *GZMB* | | | | | | | |
|---|---|---|---|---|---|---|---|
| *IL2* | 0.067 | | | | | | |
| *IFNG* | > 0.99 | > 0.99 | | | | | |
| *IL17F* | > 0.99 | 0.080 | > 0.99 | | | | |
| *IL22* | > 0.99 | < 0.001 | 0.098 | > 0.99 | | | |
| *IL21* | 0.31 | > 0.99 | > 0.99 | 0.36 | 0.005 | | |
| *CXCL13* | 0.005 | > 0.99 | 0.62 | 0.006 | < 0.001 | > 0.99 | |
| | *GZMB* | *IL2* | *IFNG* | *IL17F* | *IL22* | *IL21* | *CXCL13* |

| *GZMB* | | | | | | | |
|---|---|---|---|---|---|---|---|
| *IL2* | 0.027 | | | | | | |
| *IFNG* | > 0.99 | 0.72 | | | | | |
| *IL17F* | > 0.99 | < 0.001 | 0.20 | | | | |
| *IL22* | > 0.99 | < 0.001 | 0.078 | > 0.99 | | | |
| *IL21* | 0.83 | > 0.99 | > 0.99 | 0.014 | 0.004 | | |
| *CXCL13* | 0.14 | > 0.99 | > 0.99 | 0.001 | < 0.001 | > 0.99 | |
| | *GZMB* | *IL2* | *IFNG* | *IL17F* | *IL22* | *IL21* | *CXCL13* |

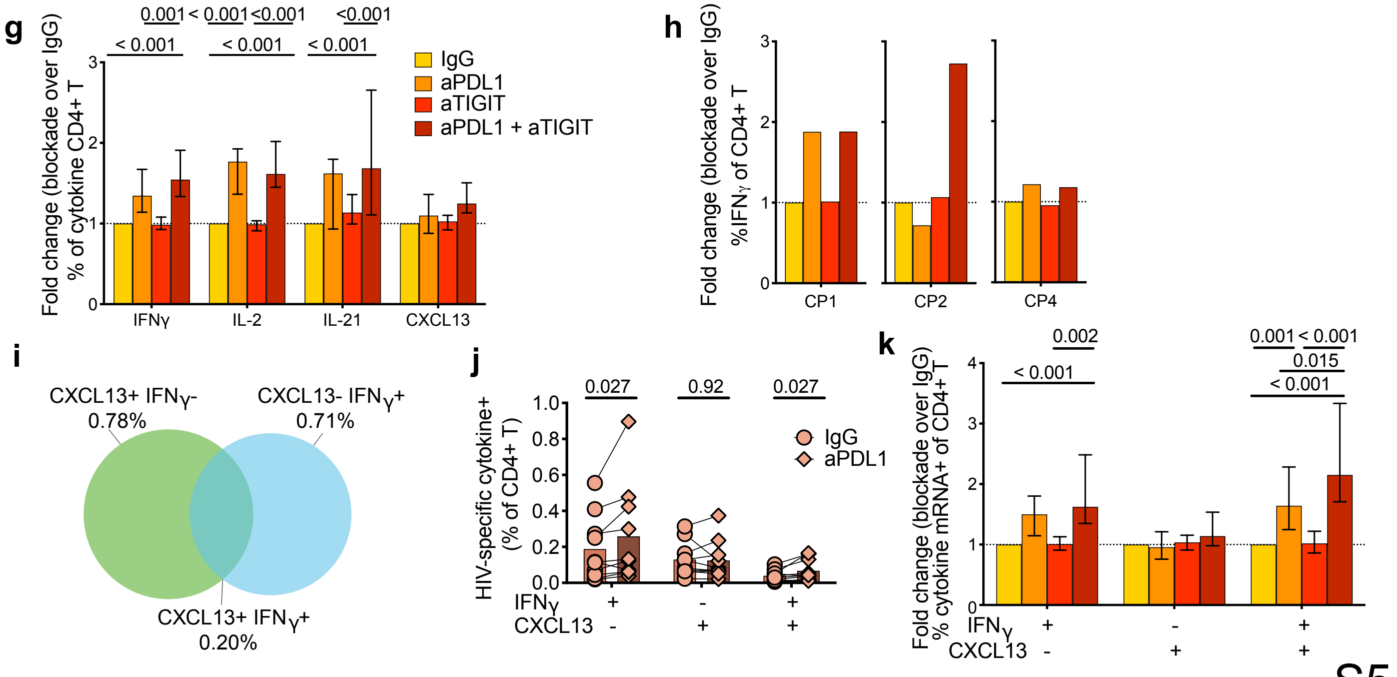


S5

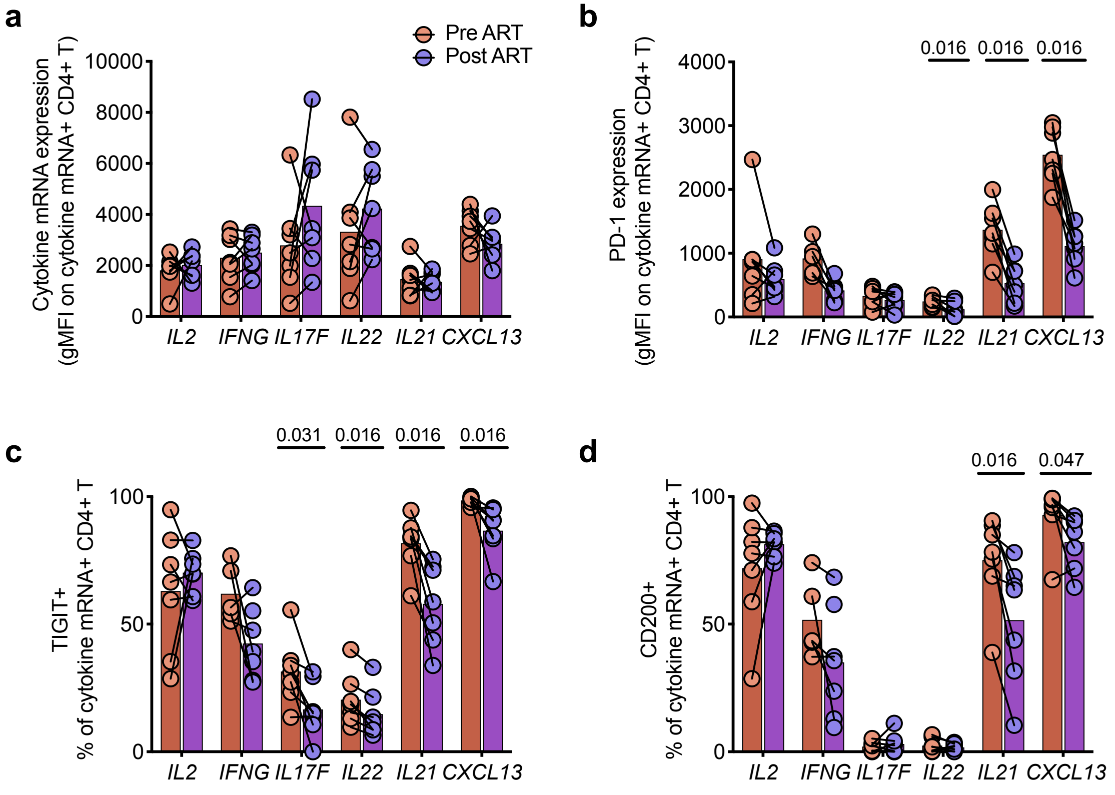
a
Pre ART
Post ART
Cytokine mRNA expression (gMFI on cytokine mRNA+ CD4+ T)
10000
8000
6000
4000
2000
0
IL2 IFNG IL17F IL22 IL21 CXCL13
b
0.016 0.016 0.016
PD-1 expression (gMFI on cytokine mRNA+ CD4+ T)
4000
3000
2000
1000
0
IL2 IFNG IL17F IL22 IL21 CXCL13
c
0.031 0.016 0.016 0.016
TIGIT+ % of cytokine mRNA+ CD4+ T
100
50
0
IL2 IFNG IL17F IL22 IL21 CXCL13
d
0.016 0.047
CD200+ % of cytokine mRNA+ CD4+ T
100
50
0
IL2 IFNG IL17F IL22 IL21 CXCL13

S6

**Supplementary table 1: Subject characteristics**

| Characteristic | CP | EC | ART |
|---|---|---|---|
| Number of subjects | 13 | 9 | 7 |
| Median (IQR) Viral load (copies/ml) | 19067 | < 40 | < 40 |
| | (6235 - 1 000 000) | (< 40 - 49) | (< 20 - 44) |
| Median (IQR) CD4 Count (cells/ul) | 329 | 591 | 640 |
| | (138 - 1036) | (369 - 744) | (361-940) |
| Median (IQR) documented years of HIV infection | 8.5 | 16.4 | 9.2 |
| | (0.25 - 23.9) | (1.2 - 28.5) | (0.9 - 24.9) |
| Median (IQR) age (years) | 41 | 46 | 43 |
| | (22 - 51) | (33 - 59) | (37 - 52) |
| Gender | | | |
| Number of Males (%) | 11 (85%) | 4 (44%) | 6 (86%) |
| Number of Females (%) | 2 (15%) | 5 (56%) | 1 (14%) |

### Supplementary Table 2: Flow cytometry panel used in AIM assay

| Antigen/Reagent | Fluorochrome | Clone | Manufacturer | Cat # | Volume/test (ul)* |
|---|---|---|---|---|---|
| Brilliant Stain buffer | - | - | BD | 563794 | 50 |
| LIVE/DEAD | Aquavivid | - | eBioscience | L34966 | 1 |
| CD14 | V500 | M5E2 | BD | 561391 | 2 |
| CD19 | V500 | H1B19 | BD | 561121 | 2 |
| CD8 | V500 | SK1 | BD | 561618 | 4 |
| CD3 | BUV395 | UCHT1 | BD | 563546 | 6 |
| CD4 | BUV496 | SK3 | BD | 564651 | 8 |
| CD69 | BV650 | FN05 | Biolegend | 310934 | 7.5 |
| CD154 | PE | TRAP1 | BD | 555700 | 20 |
| TIGIT | APC | MBSA43 | eBioscience | 17-9500 | 7.5 |
| PD-1 | BV421 | EH12.2H75 | Biolegend | 329920 | 6 |
| ICOS | PE-Cy7 | ISA-3 | eBioscience | 25-9948 | 10 |
| CD200 | PerCP-efluor710 | OX104 | eBioscience | 46-9200 | 7.5 |
| CD45RA | BUV737 | H1100 | BD | 564442 | 4 |
| CCR6** | APC-R700 | 11A9 | BD | 565173 | 7.5 |
| CXCR3** | BV605 | G025H7 | Biolegend | 353728 | 10 |
| CXCR5** | BB515 | RF8B2 | BD | 564624 | 10 |

* one test : 10M PBMC in 200ul staining buffer

** added in cull culture (0.5ml) 15 min prior to stimulation

**Supplementary Table 3: Variants of the Flow cytometry panels used in intra-nuclear transcription factor staining**

| | Antigen/Reagent | Fluorochrome | Clone | Manufacturer | Cat # | Volume/test (ul)* |
|---|---|---|---|---|---|---|
| | Brilliant Stain buffer | - | - | BD | 563794 | 10 |
| | LIVE/DEAD | Aquavivid | - | eBioscience | L34966 | 0.5 |
| | CD14 | BV480 | M5E2 | BD | 746304 | 1 |
| | CD19 | BV480 | HIB19 | BD | 746457 | 0.5 |
| | CD8 | BV480 | RPA-T8 | BD | 566121 | 0.5 |
| | CD4 | BUV496 | SK3 | BD | 564651 | 4 |
| | CD69 | BUV395 | FN50 | BD | 564364 | 2.5 |
| | CD154 | BV711 | 24-31 | Biolegend | 310838 | 5 |
| | TIGIT | PE-Cy7 | MBSA43 | eBioscience | 25-9500 | 2 |
| | PD-1 | BV605 | EH12.2H7 | Biolegend | 329924 | 5 |
| | T-BET | BV421 | O4-46 | BD | 563318 | 5 |
| | CD45RA | APC-Fire 750 | HI100 | Biolegend | 304151 | 0.5 |
| | CCR7** | BB700 | 3D12 | BD | 566438 | 5 |
| | CCR6** | BUV737 | 11A9 | BD | 564377 | 0.5 |
| | CXCR3** | BV785 | G025H7 | Biolegend | 353738 | 0.5 |
| TF exhaustion | NFATc1 | AF488 | 7A6 | Biolegend | 649604 | 5 |
| | CXCR5** | PE-Dazzle 594 | J252D4 | Biolegend | 356928 | 5 |
| | TOX | PE | TXRX10 | eBioscience | 12-6502-82 | 5 |
| | TCF-1 | AF647 | 7F11A10 | Biolegend | 655204 | 5 |
| TF Pol | CXCR5** | BB515 | RF8B2 | BD | 564624 | 2 |
| | Eomes | PE-eF610 | WD1928 | eBioscience | 61-4877-42 | 5 |
| | CD200 | PE | OXS-104 | Biolegend | 329206 | 3 |
| | RORgt | AF647 | Q21-559 | BD | 563620 | 5 |
| Optimization | Helios | PerCP eFluor710 | 22F6 | eBioscience | 46-9883 | 5 |
| | BATF | eFluor660 | MBM7C7 | ThermoFisher | 50-9860-42 | 5 |
| | Bcl6 | PE | K112-91 | BD | 561522 | 5 |

### Supplementary Table 4: Flow cytometry panel used in mRNA-Flow-FISH assay

| Antigen/Reagent | Fluorochrome | Clone | Manufacturer | Cat # | Volume/test (ul)* |
|---|---|---|---|---|---|
| Brilliant Stain buffer | - | - | BD | 563794 | 25 |
| LIVE/DEAD | Efluor-506 | - | Invitrogen | 65-0866 | 0.5 |
| CD14 | BV510 | M5E2 | Biolegend | 301842 | 3 |
| CD19 | BV510 | H1B19 | Biolegend | 302242 | 3 |
| CD56 | BV510 | NCM16.2 | BD | 563041 | 0.5 |
| CD3 | BB700 | HIT3a | BD | 742207 | 0.5 |
| CD4 | BUV496 | SK3 | BD | 564651 | 4 |
| CD8 | PE-efluor610 | RPA-T8 | eBioscience | 61-0088 | 0.5 |
| CD69 | BUV395 | FN50 | BD | 564364 | 2.5 |
| TIGIT | PE-Cy7 | MBSA43 | eBioscience | 25-9500 | 5 |
| PD1 | BV711 | EH12.2H7 | Biolegend | 329928 | 5 |
| CD200 | PE | OX-104 | Biolegend | 329206 | 3 |
| CCR6** | BUV737 | 11A9 | BD | 564377 | 2 |
| CXCR3** | BV421 | G025H7 | Biolegend | 353716 | 1 |
| CXCR5** | BV605 | J252D4 | Biolegend | 356929 | 2 |

* one test : 5M PBMC in 100ul staining buffer
** added in cull culture (0.5ml) 15 min prior to stimulation

**Supplementary Table 5: Probe panel combination used in mRNA-Flow-FISH assay**

| | Proble | Fluoro. | Manufacturer | Cat # | Volume/test (ul)* |
|---|---|---|---|---|---|
| Probe combinations | Antiviral panel | | | | |
| | *GZMB* mRNA | Type 1 | ThermoFisher | VA1-3084452 | 5 |
| | *IL2* mRNA | Type 4 | ThermoFisher | VA4-14454 | 5 |
| | *IFNG* mRNA | Type 6 | ThermoFisher | VA6-13121 | 5 |
| | Mucosal panel | | | | |
| | *IL22* mRNA | Type 1 | ThermoFisher | VA1-14439 | 5 |
| | *IL17F* mRNA | Type 4 | ThermoFisher | VA4-11185 | 5 |
| | *IL10* mRNA | Type 6 | ThermoFisher | VA6-13016 | 5 |
| | TFH Panel | | | | |
| | *IL21* mRNA | Type 1 | ThermoFisher | VA1-14117 | 5 |
| | *IL4* mRNA | Type 4 | ThermoFisher | VA4-3082434 | 5 |
| | *CXCL13* mRNA | Type 6 | ThermoFisher | VA6-15752 | 5 |
| | Modified Mucosal panel | | | | |
| | *IL22* mRNA | Type 1 | ThermoFisher | VA1-14439 | 5 |
| | *IL17F* mRNA | Type 4 | ThermoFisher | VA4-11185 | 5 |
| | *IFNG* mRNA | Type 6 | ThermoFisher | VA6-13121 | 5 |
| | Modified TFH Panel | | | | |
| | *IL21* mRNA | Type 1 | ThermoFisher | VA1-14117 | 5 |
| | *IL2* mRNA | Type 4 | ThermoFisher | VA4-14454 | 5 |
| | *CXCL13* mRNA | Type 6 | ThermoFisher | VA6-15752 | 5 |

* one test : 5M PBMC in 100ul staining buffer

**Supplementary Table 6: Flow cytometry panel used in delayed intracellular cytokine staining (d-ICS)**

| Antigen/Reagent | Fluorochrome | Clone | Manufacturer | Cat # | Volume/test (ul)* |
|---|---|---|---|---|---|
| Brilliant Stain buffer | - | - | BD | 563794 | 25 |
| LIVE/DEAD | Aquavivid | - | eBioscience | L34957 | 0.5 |
| CD14 | BV510 | M5E2 | Biolegend | 301842 | 3 |
| CD19 | BV510 | H1B19 | Biolegend | 302242 | 3 |
| CD56 | BV510 | NCAM16.2 | BD | 563041 | 0.5 |
| CD8 | BV711 | RPA-T8 | BioLegend | 301044 | 2 |
| CD69 | BUV395 | FN50 | BD | 564364 | 2.5 |
| CD4 | BUV496 | SK3 | BD | 564651 | 4 |
| CCR7** | BV650 | G043H7 | Biolegend | 353234 | 5 |
| CD45RA | APC-Fire750 | HI100 | Biolegend | 304151 | 1 |
| CXCR5** | BV421 | J252D4 | Biolegend | 356920 | 2.5 |
| IFNγ*** | PE-Cy7 | B27 | BD | 557643 | 4 |
| IL-2*** | PE-Dazzle594 | MQ1-17H12 | Biolegend | 500344 | 3.5 |
| TNFa*** | AF488 | MAb11 | Biolegend | 502915 | 2 |
| IL-21*** | eFluor660 | eBio3A3-n2 | eBioscience | 50-7219-42 | 5 |
| CXCL13*** | PE | 53610 | R&D | IC801P | 10 |

* one test : 5M PBMC in 100ul staining buffer

** added in cell culture (0.5ml) 15 min prior to stimulation

*** Stained for intracellularly
